# Ploidy shapes gemcitabine response through altered potency and delayed cell death

**DOI:** 10.64898/2026.06.01.729318

**Authors:** Vural Tagal, Jackson Cole, Tao Li, Rikhil Kumar, Jordan Albrecht, Pujan Shrestha, Didem Ilter, Daria Miroshnychenko, Stanislav Drapela, Kayode Olumoyin, John Kyei, Mark Davies, Andriy Marusyk, Stuart Maudsley, Xiaoqing Yu, Ana P. Gomes, Issam El-Naqa, Derek R. Duckett, Hyo S. Han, Noemi Andor

**Affiliations:** Department of Integrated Mathematical Oncology, H. Lee Moffitt Cancer Center & Research Institute, Tampa, FL, USA; Department of Machine Learning, H. Lee Moffitt Cancer Center & Research Institute, Tampa, FL, USA; Department of Biomedical Engineering, Texas A&M University, College Station, TX, USA; Department of Biostatistics and Bioinformatics, H. Lee Moffitt Cancer Center & Research Institute, Tampa, FL, USA; Department of Tumor Microenvironment and Metastasis, H. Lee Moffitt Cancer Center & Research Institute, Tampa, FL, USA; Velindre Cancer Center, Department of Genetics and Oncology, Cardiff University, Cardiff, UK; Department of Drug Discovery, H. Lee Moffitt Cancer Center & Research Institute, Tampa, FL, USA; Department of Breast Oncology, H. Lee Moffitt Cancer Center & Research Institute, Tampa, FL, USA

## Abstract

Aberrant tumor ploidy is a near-universal hallmark of cancer and increasingly recognized as a determinant of therapeutic response, but the mechanisms by which ploidy shapes sensitivity to specific cytotoxic agents remain unclear. Here, we investigated the relationship between ploidy and therapeutic response using pharmacogenomic reanalysis, isogenic cancer cell systems, live-cell imaging, intracellular pharmacokinetic/ pharmacodynamic (PK/PD) measurements, and mathematical modeling. Across public pharmacogenomic datasets, gemcitabine emerged as a low-ploidy-selective cytotoxic agent. In matched isogenic low- and high-ploidy cell systems, higher-ploidy cells were consistently less sensitive to gemcitabine across multiple lineages. Live-cell imaging and PK/PD measurements in near-diploid and near-tetraploid SUM-159 cells showed that both states formed intracellular dFdCTP, active form of gemcitabine; but, high-ploidy cells exhibited weaker and slower treatment responses, with delayed accumulation of cell death. To quantify these differences, we developed a delay-aware live/dead model driven by intracellular dFdCTP exposure. The model identified both reduced effective gemcitabine potency and a substantially longer delay from intracellular drug action to observed death in high-ploidy cells (17.5 hours in near-diploid cells versus 42.5 hours in near-tetraploid cells). Interpreting these fitted quantities alongside checkpoint signaling and metabolomic profiling suggests that high-ploidy cells convert intracellular gemcitabine exposure less efficiently into replication-stress signaling, nucleotide-metabolic disruption, and cytotoxic commitment. Together, these results establish ploidy as a determinant of both the magnitude and timing of gemcitabine response and provide a quantitative framework for linking intracellular drug exposure to delayed cytotoxic outcomes across ploidy states.

## Introduction

Chromosomal instability (CIN) and the resultant aneuploidy have long been recognized as a hallmark of cancer. Aneuploidy occurs in 90% of solid tumors and a majority of hematologic malignancies, while 36% of cancers harbor evidence of whole-genome doubling (WGD), with higher rates observed in metastatic disease. [1, 2]. These genome-scale alterations reshape cell proliferation patterns, gene dosage, karyotypic evolution, immune exclusion and stress tolerance while promoting more CIN, buffering deleterious chromosome alterations and creating new dependencies associated with replication, mitosis and proteostasis [1, 3, 4]. Thus, emerging evidence has shown that aneuploid and polyploid states in cancer are not merely descriptive but functionally relevant to tumorigenesis and therapy response [5–7]. Yet despite growing interest in ploidy as a biomarker of tumor evolution and therapeutic sensitivity, it remains unclear whether baseline ploidy determines the cellular response to specific anti-cancer agents.

Aneuploid and WGD-positive cancer cells differ not only in the sequence or copy number of individual genes, but also in the scale of the genome that must be duplicated, segregated, and repaired after damage. A doubled genome, therefore, imposes greater demands on DNA replication machinery, nucleotide metabolism, mitotic fidelity, protein turnover and checkpoint control while also providing increased survival fitness against DNA damage and chromosome loss. [1, 3, 4, 8–10]. This duality creates distinct biological liabilities: WGD can increase cellular burden and generate new vulnerabilities, but it can also enable tolerance of chromosome missegregation and other forms of genome damage that may be lethal in lower-ploidy cells. However, whether these genome-scale differences directly determine how cancer cells convert drug exposure into irreversible cytotoxic damage remains a central unresolved question.

One setting in which ploidy may be especially consequential is replication-stress-inducing therapies. Therapeutic agents are often interpreted as though a given extracellular concentration produces comparable intracellular stress across malignant cells. However, for drugs that perturb DNA synthesis, the effective drug pressure experienced by a cell may depend on genome size, endogenous nucleotide pools, cell-cycle state, checkpoint responsiveness and the ability to survive replication-associated chromosome damage [11–13]. Since cells with higher DNA content require larger nucleotide pools to replicate the genome, they may also be selected for enhanced metabolic buffering and tolerance of imperfect genome duplication. Conversely, lower-ploidy cells, with less chromosomal redundancy, may be more vulnerable when replication stress is converted into chromosome missegregation, micronuclei or lethal post-mitotic imbalance [3, 6, 7, 14, 15]. Thus, because whole-genome doubling changes both DNA content and the tolerance landscape for chromosome missegregation, ploidy has the capacity to influence not only how much damage a drug causes; but also the therapeutic response at multiple levels including intracellular drug activation, competition with endogenous nucleotide pools, replication-stress signaling, checkpoint activation, mitotic failure and the timing of cell death.

Gemcitabine acts as a nucleoside analogue that inhibits ribonucleotide reductase, depletes dNTP pools and induces severe DNA replication stress [19, 20]. Established mechanisms of gemcitabine resistance include reduced intracellular drug export or activation through low hENT1/SLC29A1 or dCK [21–23]; increased drug inactivation through CDA or NT5C1A [24, 25]; and metabolic rescue through upregulation of RRM1/RRM2, which restores dNTP availability and blunts gemcitabine-induced fork collapse [26–28]. In parallel, resistant cells have been demonstrated to tolerate replication stress through stronger checkpoint and fork-protection programs, including ATR/CHK1/WEE1-, NEK9-, or CK2-dependent buffering, thereby delaying or preventing conversion of S-phase damage into lethal mitosis [29–32]. These canonical mechanisms primarily explain variation in intracellular drug burden, fork collapse, and checkpoint control; however, whether the same intracellular active-metabolite exposure generates equivalent pharmacodynamic consequences across different ploidy states remains completely unknown.

In this study, we integrated pharmacogenomic analysis, matched isogenic cancer-cell systems, intracellular pharmacokinetic/ pharmacodynamic measurements (PK/PD), live-cell imaging, metabolomics, checkpoint signaling, in vivo mouse models and mathematical modeling to test how ploidy shapes therapeutic response. Across public drug-response datasets, gemcitabine emerged as a candidate low-ploidy-selective cytotoxic agent. In experimentally controlled low- and high-ploidy isogenic systems, high-ploidy derivatives were consistently less sensitive to gemcitabine across multiple cancer lineages. In matched near-diploid and near-tetraploid cancer cells, both ploidy states converted gemcitabine into its active intracellular form, dFdCTP, yet high-ploidy cells showed weaker checkpoint activation, milder nucleotide-metabolic perturbation, reduced effective cytostatic and cytotoxic potency, and delayed accumulation of cell death. Together, our findings support a model in which ploidy quantitatively shapes gemcitabine pharmacodynamics by modifying how intracellular gemcitabine exposure is translated into nucleotide stress, checkpoint signaling and cytotoxicity. This work positions ploidy as a functional variable in chemotherapy response and suggests that DNA-content state has to be considered when interpreting, modeling and eventually exploiting drug sensitivity in cancer.

## Results

### Ploidy stratifies cytotoxic and targeted therapy-response patterns across cancers

To determine whether ploidy is associated with distinct therapeutic response programs, we integrated drug-sensitivity measurements from the Genomics of Drug Sensitivity in Cancer (GDSC) database [33] with cell line ploidy estimates (Cell Line Passports database) [34] (Van Der Meer, D., et al. Nucleic Acid Research, 2019) (Fig. 1A). Classified compounds were grouped into broad therapeutic classes, and enrichment of ploidy-associated drugs were evaluated within each cancer type and all cancers overall (Methods 1.3). Across cancers, drug-class enrichments resolved into low-ploidy-biased, high-ploidy-biased, and bidirectional/context-dependent patterns (Fig. 1B,C; Supplementary Fig. 1A-B). Chemotherapy agents showed the strongest low-ploidy bias, with significant enrichment in seven low-ploidy contexts (All cancers, BRCA, ALL, BLCA, COREAD, HNSC, unclassfied) compared with one high-ploidy context (*q ≤* 0.05)(Fig. 1B). Kinase inhibitors were significantly enriched in both directions, but were more frequently associated with high-ploidy sensitivity occurring in four high-ploidy cancer contexts (All cancers, BRCA, COREAD, GBM) compared to two low-ploidy contexts (SCLC, DLBC), consistent with a context-dependent but high-ploidy-skewed pattern (Fig. 1C). By contrast, other categories were not directionally uniform across cancer contexts (Fig. 1B,C; Supplementary Fig. 1A-B). These findings indicate that ploidy broadly stratifies therapeutic response, with cytotoxic agents preferentially associated with low-ploidy sensitivity and targeted agents displaying high-ploidy dependence.

**Figure 1.**
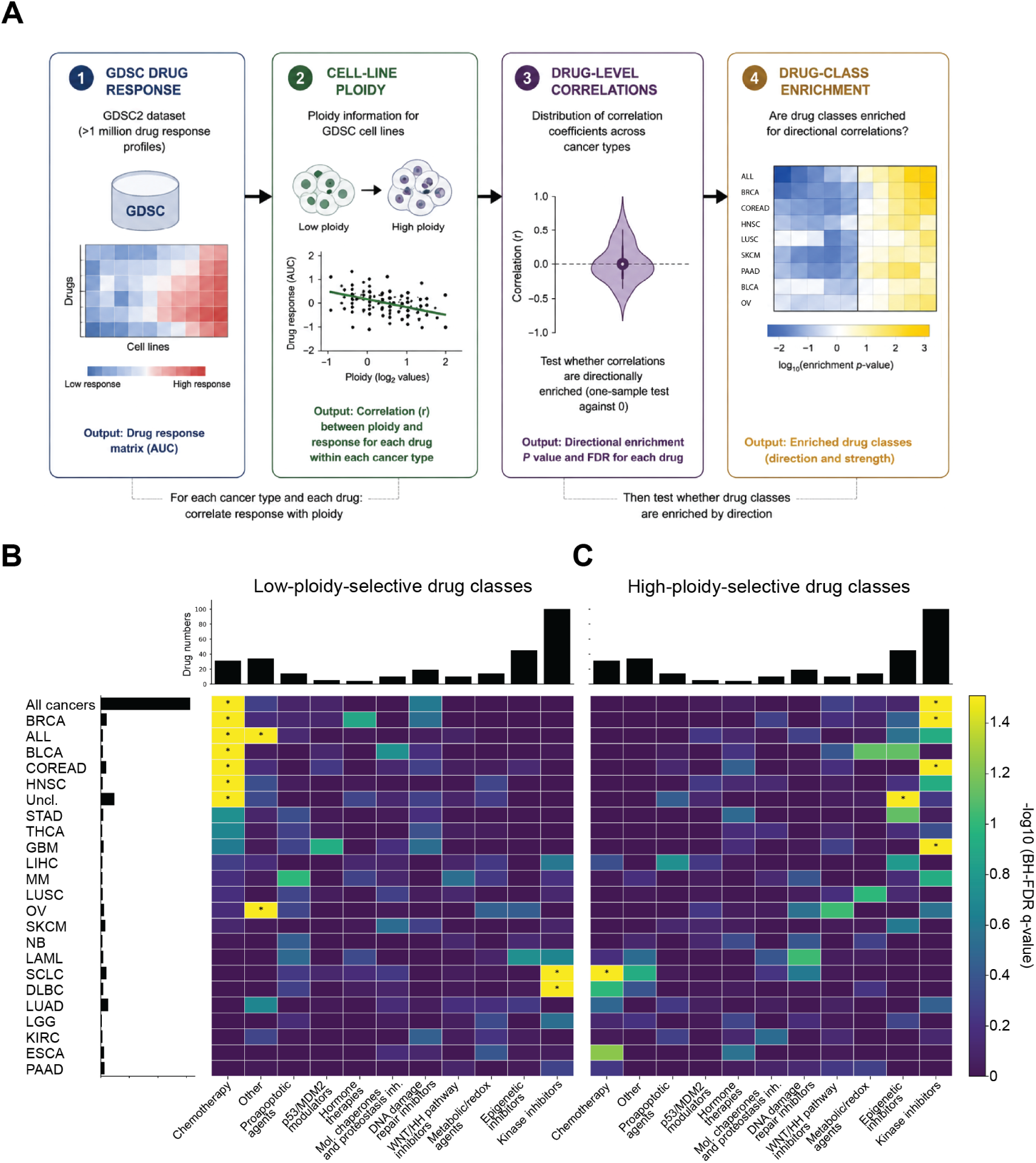
Ploidy-associated drug-class enrichment across cancer cell lines. **(A)** Workflow schematic. GDSC drug-response measurements were integrated with cell-line ploidy estimates. For each cancer type and drug, drug response across matched cell lines was correlated with ploidy, and drugs were then stratified by the direction of the ploidy–response association before testing for drug-class enrichment. **(B)** Heatmap showing drug classes enriched among compounds with response patterns consistent with preferential low-ploidy sensitivity. Rows indicate cancer types and columns indicate curated drug classes. Color represents enrichment significance as *−* log_10_(*q − value*), with larger values indicating stronger enrichment. **(C)** Heatmap showing drug classes enriched among compounds with response patterns consistent with preferential high-ploidy sensitivity, using the same row order, column order, and color scale as in panel B. Stars indicate nominal enrichment significance at *q ≤* 0.05. Exact zero values from permutation testing were plotted at the empirical permutation floor to avoid infinite *−* log_10_(*q − value*) values. (GDSC, Genomics of Drug Sensitivity in Cancer; AUC, area under curve; BRCA, breast invasive carcinoma; ACC, adrenocortical carcinoma; ALL, acute Lymphoblastic Leukemia; BLCA, bladder urothelial carcinoma; COREAD, colorectal adenocarcinoma; HNSC, head and neck squamous cell carcinoma; STAD, stomach adenocarcinoma; THCA, thyroid carcinoma; GBM, glioblastoma multiforme; LIHC, liver hepatocellular carcinoma; MM, multiple myeloma; LUSC, lung squamous cell carcinoma; OV, ovarian serous cystadenocarcinoma; SKCM, skin cutaneous melanoma; NB, neuroblastoma; LAML, Acute Myeloid Leukemia; SCLC, small cell lung carcinoma; DLBC, lymphoid neoplasm diffuse large B-cell carcinoma; LUAD, lung adenocarcinoma; LGG, brain lower grade glioma; KIRC, kidney renal clear cell carcinoma; ESCA, oesophageal carcinoma; PAAD, pancreatic adenocarcinoma.)

### Pharmacogenomic analyses and isogenic cell systems nominate gemcitabine as a low-ploidy selective agent in breast cancer

Next, we asked whether individual cytotoxic agents exhibited reproducible ploidy-associated response patterns across independent pharmacogenomic resources. Gemcitabine emerged as a leading candidate ranking among the top hits across three independent pharmacogenomic datasets. In breast cancer cell lines from CCLE [16,17], higher ploidy was associated with reduced gemcitabine sensitivity (Fig. 2A). Concordant associations were observed in independent analyses of GDSC breast cancer models and NCI-60-derived response data for standard anticancer agents [33] [18]. (Fig. 2B,C). The convergence of these orthogonal datasets identified gemcitabine as a reproducible low-ploidy-selective cytotoxic agent and provided the rationale for direct testing in controlled and lineage-matched isogenic ploidy systems.

**Figure 2.**
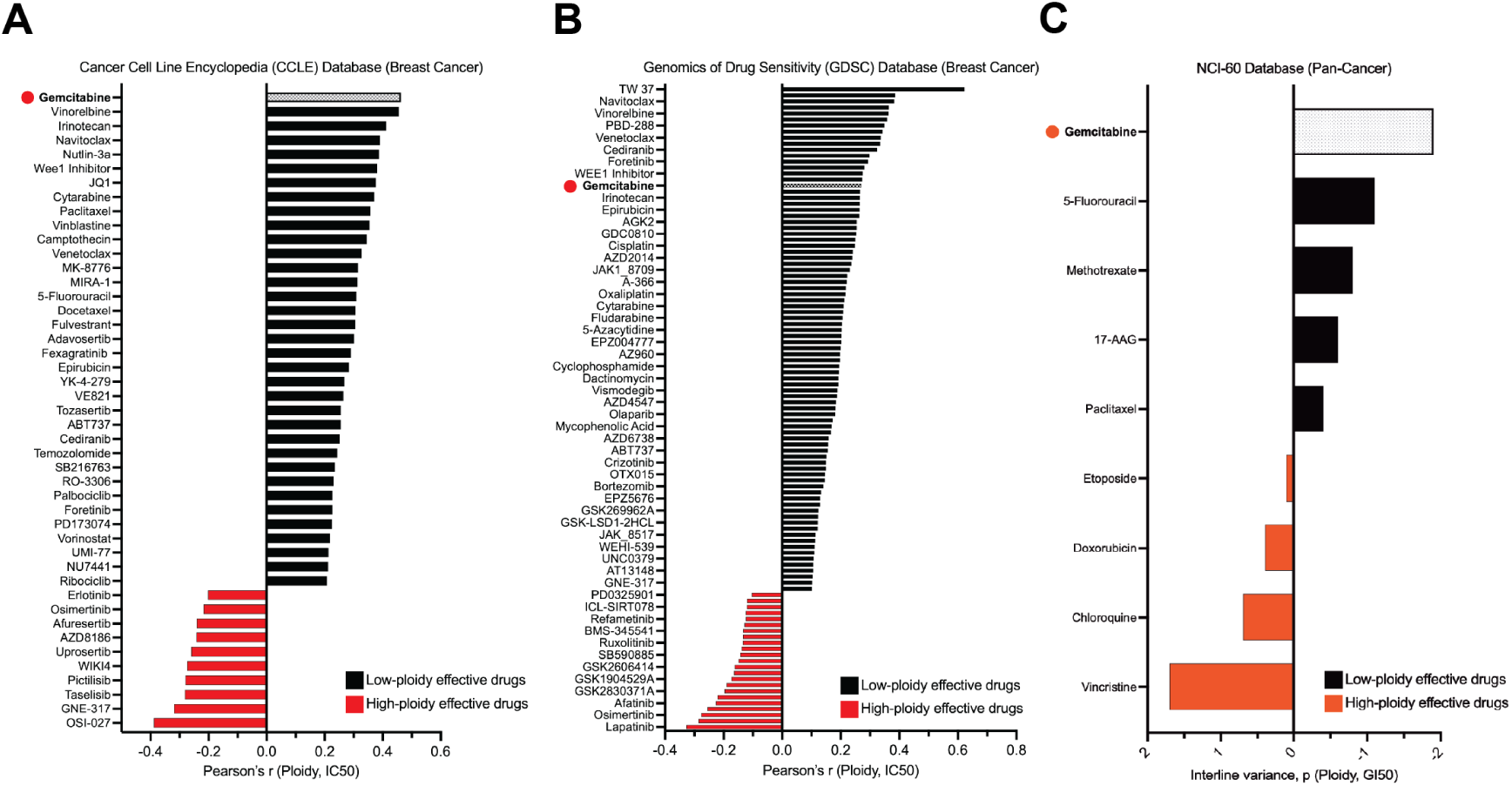
Ploidy predicts gemcitabine response across independent pharmacogenomic datasets. (A) Local reproduction of the CCLE breast cancer ploidy–drug-response analysis [16], using matched cell-line ploidy estimates and drug-resistance Z-score across 26 breast cancer cell lines. Positive correlations indicate higher drug-resistance z-score in higher-ploidy lines, consistent with relatively greater sensitivity in lower-ploidy lines. Data source: CCLE [17]. (B) Pearson correlation between cell-line ploidy and GDSC drug-response z-score is shown for drugs administered to breast invasive carcinoma (BRCA). (C) Independent pharmacogenomic data nominate gemcitabine as low-ploidy selective. In the NCI-60-derived analysis of standard cancer therapies, gemcitabine showed the strongest negative correlation between ploidy and drug sensitivity among the listed standard agents, consistent with preferential sensitivity of lower-ploidy cells [18].

To test whether the pharmacogenomic association persisted within matched genetic backgrounds, we generated a panel of six isogenic lineage pairs derived from four cancer cell lines. The panel included low-ploidy states ranging from 2N to 4.2N and matched high-ploidy derivatives ranging from 4N to 6.5N, with ploidy differences spanning 0.1–2.6 haploid genome equivalents (Fig. 3A–C). Because the high-ploidy states were engineered through independent routes of ploidy alteration, this design reduced confounding by lineage background and by the mechanism through which increased ploidy was acquired.

**Figure 3.**
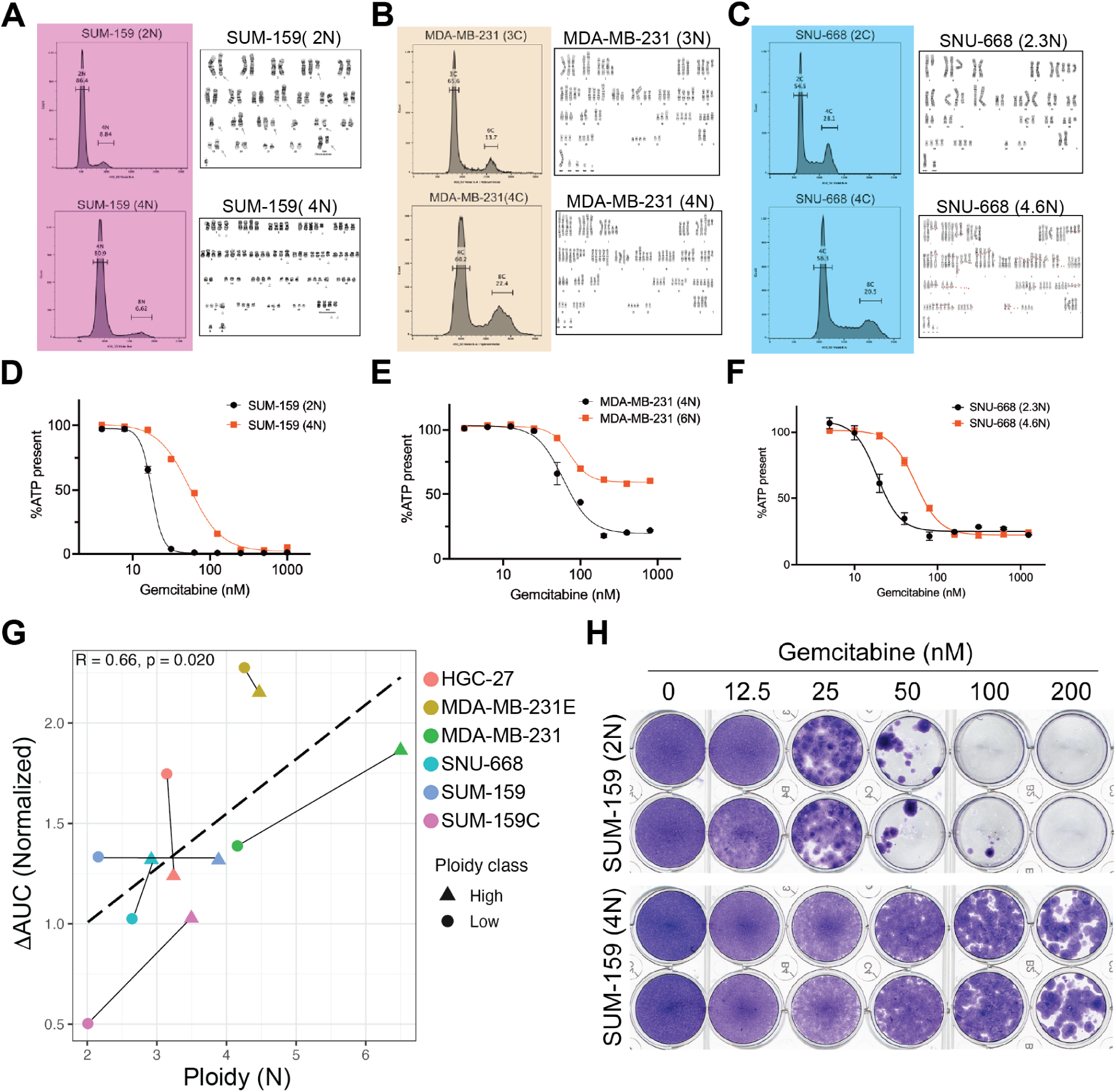
High-ploidy cells are gemcitabine-resistant. (**A**) Flow-cytometric DNA-content profiles and representative G-banded karyotypes of matched low- and high-ploidy derivatives of SUM-159, (**B**) MDA-MB-231 and (**C**) SNU-668 lines. (**D**) Gemcitabine dose-response curves for SUM-159, (**E**)MDA-MB-231 and (**F**) SNU-668 isogenic pairs. Cells were treated with gemcitabine at the concentrations shown and cell viability was measured with a CellTiter-Glo assay, which measures cellular ATP as an indicator of living cells. (**G**) Pearson correlation between mean cell ploidy and normalized gemcitabine dose-response AUC across individual samples from the isogenic panel. Higher AUC indicates greater residual growth or viability and, consequently, lower gemcitabine sensitivity (*n* = 12, *r* = 0.657, *p* = 0.020). (**H**) SUM-159 cells were treated with serial concentrations of gemcitabine for seven days and cell growth quantified with a crystal violet staining assay.

Across isogenic SUM-159, MDA-MB-231 and SNU-668 cell line models, high-ploidy derivatives consistently retained greater viability following gemcitabine treatment than their lower-ploidy counterparts (Fig. 3D–F). Across individual samples in the isogenic cell line panel, mean ploidy correlated positively with normalized gemcitabine dose response in the means of area under curve (AUC), indicating progressively greater residual growth or viability and lower gemcitabine sensitivity with increasing ploidy (Fig. 3G) (Pearson *r* = 0.657, *p* = 0.020). Furthermore, pairwise comparisons showed the same directional relationship as the the two largest increases in AUC occurred in the isogenic pairs with largest increases in ploidy (Fig. 3G). In an orthogonal assay for cell growth, SUM-159 cells were stained with crystal violet after seven days treatment with gemcitabine and cell densities were compared with the density of cells treated with vehicle. As summarized in Fig. 3H, we observed that growth of SUM-159 cells was inhibited at lower concentrations of gemcitabine in near-diploid cells whereas near-tetraploid SUM-159 cells were more tolerant to the treatment.

### dFdCTP accumulation leads to checkpoint activation and cytotoxicity in a ploidy-dependent manner

To investigate the basis for the differential cytotoxicity resulting from gemcitabine treatment in low and high-ploidy breast cancer cells, we focused mechanistic analyses on the SUM-159 pair because it provided a substantial ploidy separation with a triple-negative breast cancer background. We first explored whether the reduced sensitivity of near-tetraploid (4N) SUM-159 cells (henceforth SUM-159 (4N) cells) reflected lower intracellular activation of gemcitabine. Gemcitabine requires intracellular activation to phosphorylated metabolites including dFdCDP and dFdCTP. These metabolites disrupt DNA synthesis through complementary mechanisms: dFdCDP inhibits ribonucleotide reductase and perturbs deoxynucleotide availability while dFdCTP competes with endogenous cytidine nucleotides and can be incorporated into DNA, promoting replication stress and delayed cytotoxicity (Fig. 4A). To distinguish drug exposure from downstream response, we quantified intracellular gemcitabine metabolites over time and compared these trajectories with cell fate. Time-resolved intracellular PK/PD profiling showed that both ploidy states generated active gemcitabine metabolites. Unexpectedly, SUM-159 (4N) cells accumulated more dFdCTP, active form of gemcitabine, than near-diploid (2N) SUM-159 cells (henceforth SUM-159 (2N) cells) whereas the dFdU (inactive form) trajectories were broadly similar between the two states (Fig. 4B-C). Despite greater dFdCTP accumulation, our live-cell imaging observations demonstrated that SUM-159 (4N) cells exhibited delayed and reduced cytotoxicity in comparison to SUM-159 (2N) cells, and retained substantially more viable cells following the treatment (Fig. 4D). Moreover, gemcitabine-treated SUM-159 (4N) cells also developed enlarged nuclei and cell size, consistent with accumulation in S or G2/M phases of cell cycle (Fig. 4D). Overall, our findings suggested that gemcitabine treatment might induce ploidy-dependent replication stalling leading to differential DNA damage and ultimately cell death.

**Figure 4.**
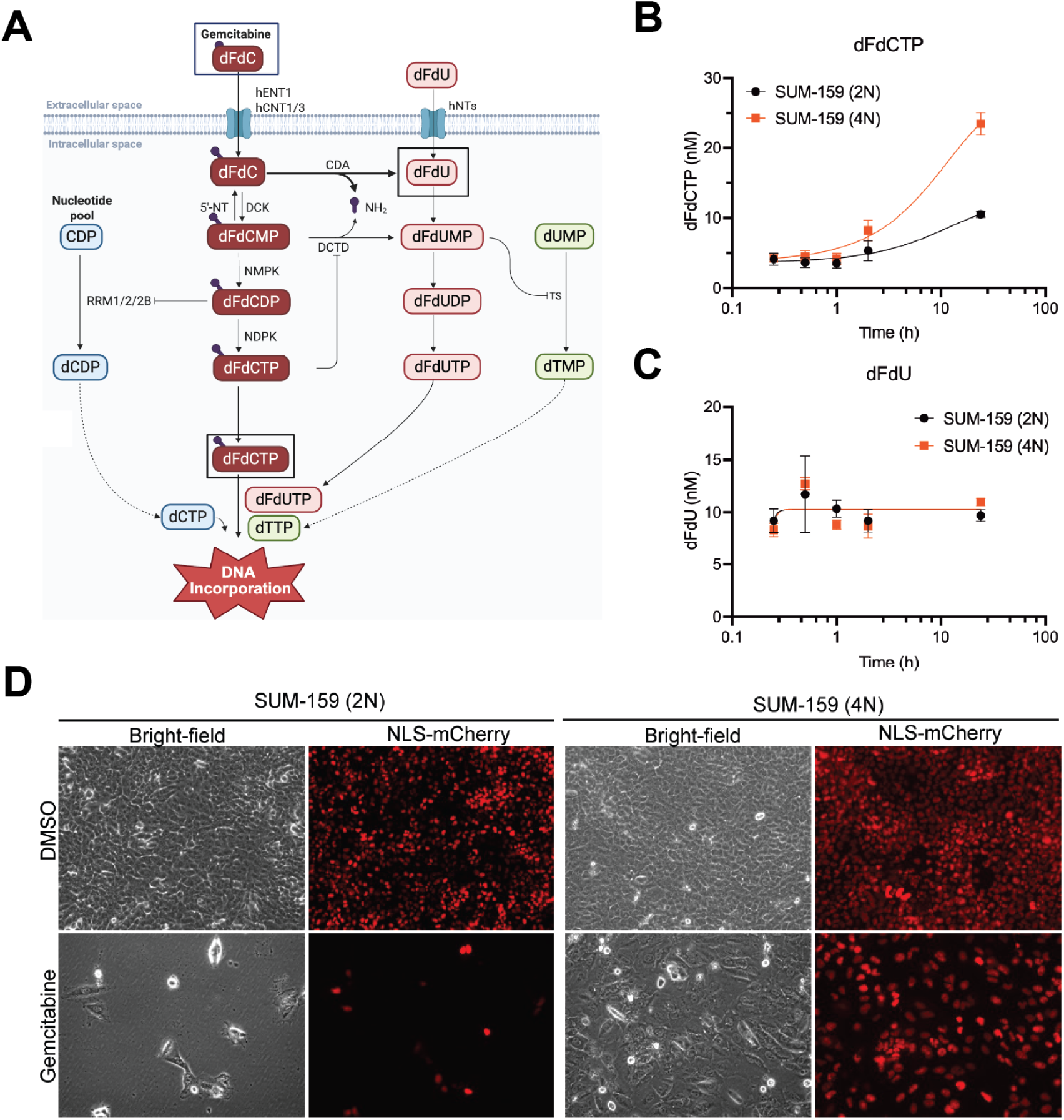
Gemcitabine metabolism, intracellular dFdCTP accumulation and ploidy-dependent response in SUM-159 cells. **(A)** Schematic of gemcitabine (dFdC) uptake and metabolism. Extracellular gemcitabine (dFdC) enters cells and is sequentially phosphorylated through the deoxycytidine salvage pathway to dFdCMP, dFdCDP and dFdCTP, the active triphosphate metabolite that can be incorporated into DNA. Gemcitabine metabolites can also be deaminated to dFdU species, and dFdCDP inhibits ribonucleotide reductase, thereby perturbing endogenous deoxynucleotide synthesis. **(B)** Time-resolved intracellular measurements of dFdCTP and **(C)** dFdU in matched SUM-159 near-diploid (2N) and near-tetraploid (4N) cells following gemcitabine exposure. Near-tetraploid cells accumulated more dFdCTP, whereas dFdU concentrations were broadly comparable between 2N and 4N cells. **(D)** Representative bright-field and NLS-mCherry images of SUM-159 2N and 4N cells four days after treatment with DMSO or gemcitabine. Gemcitabine caused marked loss of viable/nuclear signal in 2N cells whereas 4N cells retained more cells and nuclear signal under the same treatment condition, consistent with relative gemcitabine resistance despite detectable active-metabolite accumulation.

Because active dFdCTP formation was not reduced in SUM-159 (4N) cells, we hypothesized that the divergence in therapeutic response between ploidy states emerged downstream, at the level of replication-stress signaling. To determine whether ploidy alters the conversion of intracellular gemcitabine exposure into downstream replication-stress signaling, we measured checkpoint activation 24 hours after gemcitabine treatment in matched SUM-159 (2N) and SUM-159 (4N) cells. Gemcitabine induced phosphorylation of CHK1, and to a lesser extent CHK2, at lower administered doses in SUM-159 (2N) cells than in SUM-159 (4N) cells (Fig. 5G). Thus, because CHK1 is a canonical effector of replication-stress signaling, these data confirmed that low-ploidy cells engaged replication-stress and DNA-damage signaling at concentrations that produced comparatively weak checkpoint activation in high-ploidy cells. Together with intracellular dFdCTP measurements and live cell imaging, these findings indicate that high-ploidy resistance reflects attenuated conversion of gemcitabine exposure into checkpoint activation and delayed cytotoxicity. SUM-159 (4N) cells generated abundant intracellular dFdCTP but converted that exposure less efficiently into checkpoint activation and cytotoxicity, indicating that resistance arises downstream of—or in parallel with—gemcitabine activation.

**Figure 5.**
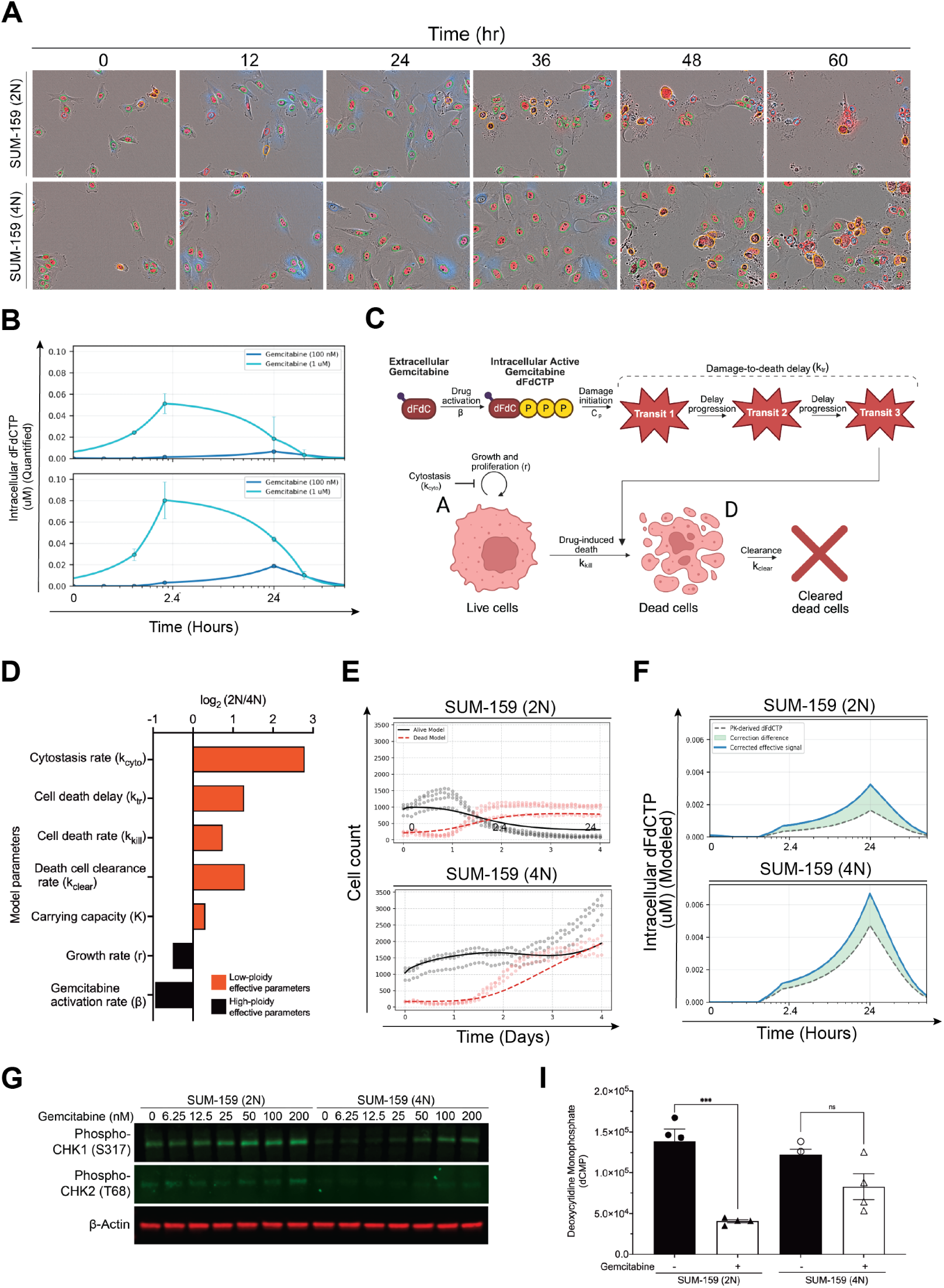
PK/PD-informed live/dead modeling of delayed gemcitabine cytotoxicity. (**A**) Live-cell imaging of gemcitabine-treated near-2N and near-4N isogenic SUM-159 cells. (**B**) Baseline-subtracted intracellular dFdCTP exposure used to inform the pharmacodynamic model. Measured dFdCTP profiles and log-linear extrapolations at the calibrated reference concentrations are illustrated. (**C**) Schematic of the delay-aware live/dead model. Intracellular dFdCTP is converted into an effective drug-action signal that suppresses live-cell growth and initiates progression through a latent transit chain representing the interval between damage initiation and observed cell death. The terminal transit compartment drives conversion of live cells to dead cells, which subsequently undergo clearance. (**D**) Ploidy-associated differences in fitted model parameters. Bars show log_2_-transformed parameter fold changes comparing 2N with 4N cells. Positive values indicate lower fitted values in 2N cells. The 4N fit shows a lower transit rate, corresponding to a longer inferred delay from effective dFdCTP exposure to death commitment. (**E**) Representative observed and model-predicted live- and dead-cell trajectories for near-diploid and near-tetraploid SUM-159 cells treated with 25 nmol/L gemcitabine. Points denote observed object counts, and curves denote the corresponding model predictions. (**F**) Baseline-subtracted intracellular dFdCTP exposure functions used to inform the pharmacodynamic model. Dose-calibrated dFdCTP-derived signal surfaces across the modeled gemcitabine concentration range are illustrated for 25 nM gemcitabine. (**G**) Immunoblot analysis of phosphorylated CHK1 and CHK2 24 hours after gemcitabine treatment in 2N and 4N SUM-159 cells. *β*-actin served as a loading control. (**I**) Metabolomic abundance of deoxycytidine monophosphate (dCMP) in untreated and gemcitabine-treated near-diploid and near-tetraploid SUM-159 cells. Gemcitabine produced a larger reduction in dCMP abundance in the near-diploid state. *** p = 0.0003, ns indicates “not significant”.

### PK/PD-informed model quantifies ploidy-dependent gemcitabine potency and cell death latency

To quantify how ploidy altered the relationship between intracellular exposure and cell fate, we developed a delay-aware live/dead compartment model (Fig. 5C) in which intracellular dFdCTP produces immediate cytostasis and feeds a latent transit process, culminating in cell death (Fig. 5C; Supplementary methods 2.1; Supplementary Table 2). The model was informed by time-resolved dFdCTP measurements and jointly calibrated to high-temporal-resolution live-cell imaging with 8,780 live- and dead-cell observations across gemcitabine concentrations, replicates, and ploidy states in matched SUM-159 (2N) and SUM-159 (4N) cells (Fig. 4E, Fig. 5A-B). Because the PK-derived signal alone did not adequately reproduce the low- and intermediate-dose responses, we included a nonlinear dose correction comprising a ploidy-specific power-law term and a shared Hill gate (Eq. (2)). The fitted gate had *EC*_50_ = 0.0185 *µ*M and a dimensionless Hill coefficient *h* = 2.69, placing the transition in effective drug action within the dose range where substantial responses emerged (Fig. 5F; Supplementary Table 3). Overall, the fitted correction suppressed effective drug action at the lowest doses, amplified the intermediate response range, and compressed higher doses relative to the reference dose (Fig. 5F; Supplementary Table 3). Here, the ploidy-specific correction terms indicate that the fitted model applies a stronger dose-scaling correction in SUM-159 (2N) than in SUM-159 (4N) cells across the low-to-intermediate dose range, although the final effective intracellular signal also depends on the measured PK-derived dFdCTP surface.

The fitted model reproduced the major features of the live and dead object trajectories across both SUM-159 (2N) and SUM-159 (4N) cells, shown for a representative 25 nM comparison in Fig. 5E and across the full dose series in Supplementary Fig. 2. In particular, the model captured the high-dose decline in live cells, the delayed accumulation of dead objects, and the qualitative separation between low-dose, intermediate-dose, and high-dose gemcitabine responses. The best-fitting model contained *n*_tr_ = 5 transit compartments, a dimensionless compartment count (Supplementary Table 1).

Effective gemcitabine potency was lower in SUM-159 (4N) cells. With the effective dFdCTP signal expressed in *µ*M-equivalent units, fitted cytotoxic potency was *k*_kill,2*N*_ = 217.3 day*^−^*^1^ *µ*M*^−^*^1^ versus *k*_kill,4*N*_ = 129.9 day*^−^*^1^ *µ*M*^−^*^1^, approximately 1.7-fold higher in 2N cells. Fitted cytostatic potency was *k*_cyto,2*N*_ = 8041 *µ*M*^−^*^1^ versus *k*_cyto,4*N*_ = 1163 *µ*M*^−^*^1^, approximately 6.9-fold higher in 2N cells. Thus, at a comparable effective intracellular signal, the model attributed stronger proliferation suppression and cell killing to the low-ploidy state (Fig. 5D; Supplementary Table 1). The response was also substantially delayed in SUM-159 (4N) cells. Fitted transit rates of *k*_tr,2*N*_ = 6.84 day*^−^*^1^ and *k*_tr,4*N*_ = 2.82 day*^−^*^1^ corresponded to modeled mean delays *n*_tr_*/k*_tr_ of 0.73 days (*∼* 17.5 hours) and 1.77 days (*∼* 42.5 hours), respectively. Together, the potency and delay estimates indicate that high-ploidy cells responded less strongly and died later, with an approximately 2.4-fold longer modeled delay to death (Fig. 5D; Supplementary Table 1).

Together, these results reveal that high-ploidy resistance comprises both a reduction in effective drug potency and a prolonged delay in cytotoxic execution. The model therefore defines a thresholded, nonlinear, and ploidy-dependent relationship between administered gemcitabine, intracellular dFdCTP, and eventual cell death.

### Gemcitabine induces broader nucleotide-metabolism remodeling in low-ploidy cells

To investigate whether attenuated checkpoint activation could reflect reduced upstream nucleotide-pathway perturbation rather than only decreased checkpoint sensitivity to damage, we performed untargeted metabolomics in isogenic SUM-159 (2N) and SUM-159 (4N) cells, 24 hours after gemcitabine exposure, using untreated cells as control. Principal component analysis (PCA) of metabolite intensities separated samples by both ploidy state and treatment condition, with 2N control, 2N gemcitabine-treated, 4N control, and 4N gemcitabine-treated samples forming distinct groups in principal component analysis (PCA) space (Fig. 6A). This separation indicates that both basal ploidy and gemcitabine exposure contribute to global metabolomic variation in SUM-159 cells. Importantly, the separation between control and gemcitabine-treated SUM-159 (2N) samples was more prominent than the corresponding separation in SUM-159 (4N) samples, consistent with the broader treatment-response signal detected in subsequent differential metabolite analyses. Using a threshold of at least two-fold change with statistical significance (p < 0.05), 63 metabolites were classified as strong gemcitabine-response or ploidy-dependent differential-response features. Metabolite response category analysis showed that gemcitabine induced a broader metabolic response in SUM-159 (2N) cells than in SUM-159 (4N) cells, with a larger number of 2N-specific response metabolites compared with 4N-specific responses (Fig. 6B). Specifically, this classification revealed a larger number of 2N-specific metabolic responses than 4N-specific responses, with 37 metabolites showing 2N-specific changes compared with only 15 metabolites showing 4N-specific changes. Eight metabolites demonstrated common gemcitabine-associated changes across both ploidy states while three metabolites were classified as interaction-only features. These response category counts confirm that gemcitabine triggers a broader and more pronounced metabolomic response in SUM-159 (2N) cells whereas SUM-159 (4N) cells exhibit an attenuated or buffered metabolic response.

**Figure 6.**
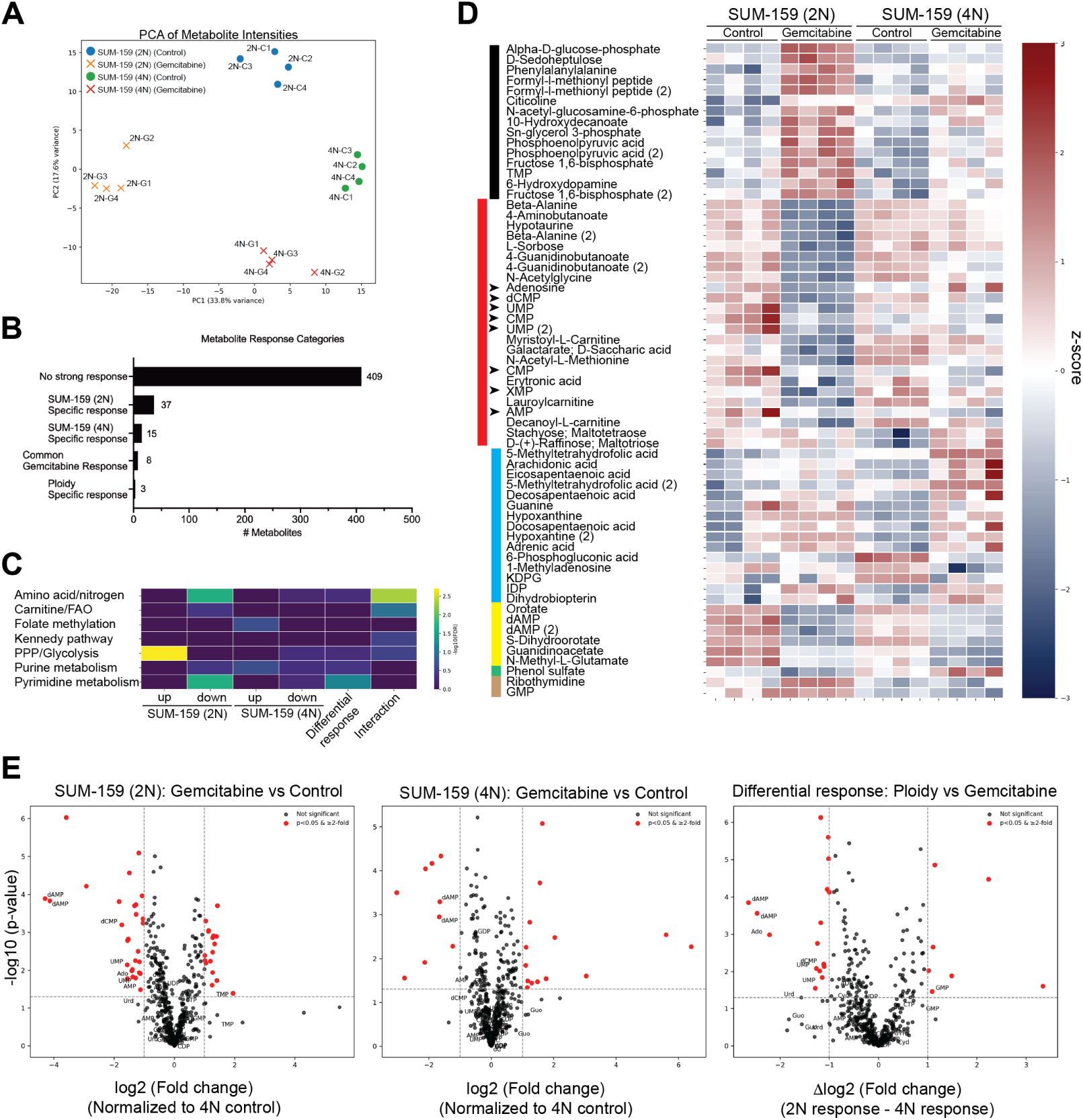
Ploidy state determines gemcitabine-induced metabolic remodeling and nucleotide depletion in SUM-159 cells. (**A**) Principal component analysis (PCA) of untargeted metabolomics profiles from SUM-159(2N) and 4N cells under control and gemcitabine-treated conditions. Each point represents an individual biological replicate. (**B**) Response category counts for metabolites with the predefined response threshold of 2-fold change and p < 0.05. Metabolites were classified as showing no strong response, 2N-specific response, 4N-specific response, common gemcitabine response or ploidy-specific differential response. (**C**) Curated pathway-class enrichment analysis of metabolite response categories. Enrichment was assessed using curated, name-based pathway classes. (**D**) Heatmap of metabolites with 2-fold or more change and p < 0.05 response threshold or showing ploidy-dependent differential response. Rows represent metabolites ordered by response class, and columns represent biological replicates from SUM-159 2N and 4N cells under control and gemcitabine-treated conditions. Values are shown as row-scaled z-scores. Changes in nucleotide-related metabolites are indicated with arrow heads. (**E**) Volcano plots showing metabolite-level responses to gemcitabine in SUM-159 2N cells, SUM-159 4N cells, and the ploidy-dependent differential response. For the 2N and 4N plots, the x-axis represents log2 fold change after gemcitabine treatment relative to control, and the y-axis represents *−* log_10_(p-value). For the differential-response plot, the x-axis represents Δlog2 fold change, calculated as the 2N response minus the 4N response. Red points indicate metabolites passing p < 0.05 and *≥* 2-fold change. Core pyrimidine and purine metabolites are labeled to highlight nucleotide-associated changes.

Curated pathway enrichment of these response classes further identified nucleotide- and nucleotide-supporting pathways as major components of the ploidy-dependent metabolic response, including pentose phosphate pathway (PPP)/glycolysis, purine metabolism and pyrimidine metabolism (Fig. 6C). Notably, metabolites decreased after gemcitabine in SUM-159 (2N) cells were enriched for pyrimidine metabolism whereas this pattern was not observed in SUM-159 (4N) cells. These findings suggested that the low-ploidy state is associated with stronger depletion or suppression of pyrimidine- and nucleotide-related metabolite pools following gemcitabine treatment compared with the high-ploidy state. The heatmap of differentially-responsive metabolites reinforced this interpretation by showing coordinated 2N-specific changes in nucleotide-related metabolites, including dCMP, UMP, CMP, AMP, XMP, IDP, dAMP, orotate, and dihydroorotate (Fig. 6D). Finally, gemcitabine produced more extensive metabolite-level changes in SUM-159 (2N) cells and identified a subset of metabolites whose treatment response differed between ploidy states. Nucleotide-associated features were among the strongest treatment and interaction signals, supporting a model in which gemcitabine imposes a more pronounced nucleotide-metabolic stress on SUM-159 (2N) cells (Fig. 6E). Overall, these analyses demonstrate that, relative to SUM-159 (4N) cells, SUM-159 (2N) cells undergo stronger gemcitabine-induced remodeling of nucleotide metabolism, with evidence consistent with decreased pyrimidine and purine-associated metabolite abundance. Therefore, this ploidy-dependent metabolic response contributes to the greater gemcitabine sensitivity of SUM-159 (2N) cells whereas SUM-159 (4N) cells are able to buffer nucleotide-metabolic disruption and better tolerate gemcitabine-induced toxicity.

### Gemcitabine induces ploidy-dependent anabolic attenuation associated with early tumor control in vivo

To determine whether the ploidy-associated difference in gemcitabine response observed in vitro was retained in vivo, we analyzed tumor growth and single-cell RNA-sequencing (scRNA-Seq)-based transcriptomic profiles from 16 orthotopic tumors established from SUM-159 (2N) or SUM-159 (4N) cells. Within each injected-origin group, four tumors received vehicle, two received 30 mg/kg gemcitabine, and two received 120 mg/kg gemcitabine (Fig. 7A-B). Because tumor-cell ploidy could change after implantation [35–37], we first used NUMBAT-derived copy-number estimates, a well-established allele-specific copy number variation (CNV) caller for scRNA-seq data to assess endpoint ploidy [38]. Of the 14,125 tumor cells with postprocessed NUMBAT estimates, 9,832 passed the final transcriptomic quality-control criteria used for all downstream analyses. This dataset comprised 4,880 cells from SUM-159 (2N)-origin tumors and 4,952 cells from SUM-159 (4N)-origin tumors, including 5,335 cells from gemcitabine-treated tumors (Supplementary Fig. 4A). Copy-number profiles revealed substantial heterogeneity both within and between tumors (Supplementary Fig. 4A, Supplementary Fig. 5A–E). To relate their endpoint states to the injected ploidy states, we compared the per-tumor endpoint estimates with project-designated, lineage-matched culture references used as proxies for the injected populations. The reference populations had mean ploidies of 2.151N and 3.947N for SUM-159 (2N) and SUM-159 (4N), respectively. By contrast, the corresponding mouse-balanced endpoint means were 2.136N for 2N-origin tumors (range of mouse means, 1.941–2.316) and 2.319N for 4N-origin tumors (range of mouse means, 2.214–2.586) (Supplementary Fig. 5E). The mean 2N–4N separation consequently contracted from 1.795N in the reference proxies to 0.183N at harvest, an 89.8% reduction. This comparison indicates substantial convergence of the 4N-derived state toward lower endpoint ploidy. Orthogonal flow-cytometry-based DNA-content measurements of endpoint material identified dominant peaks of 1.88N–2.20N across the eight 4N-origin tumors, corroborating contraction of the ploidy separation *in vivo* (Supplementary Fig. 6A,B).

**Figure 7.**
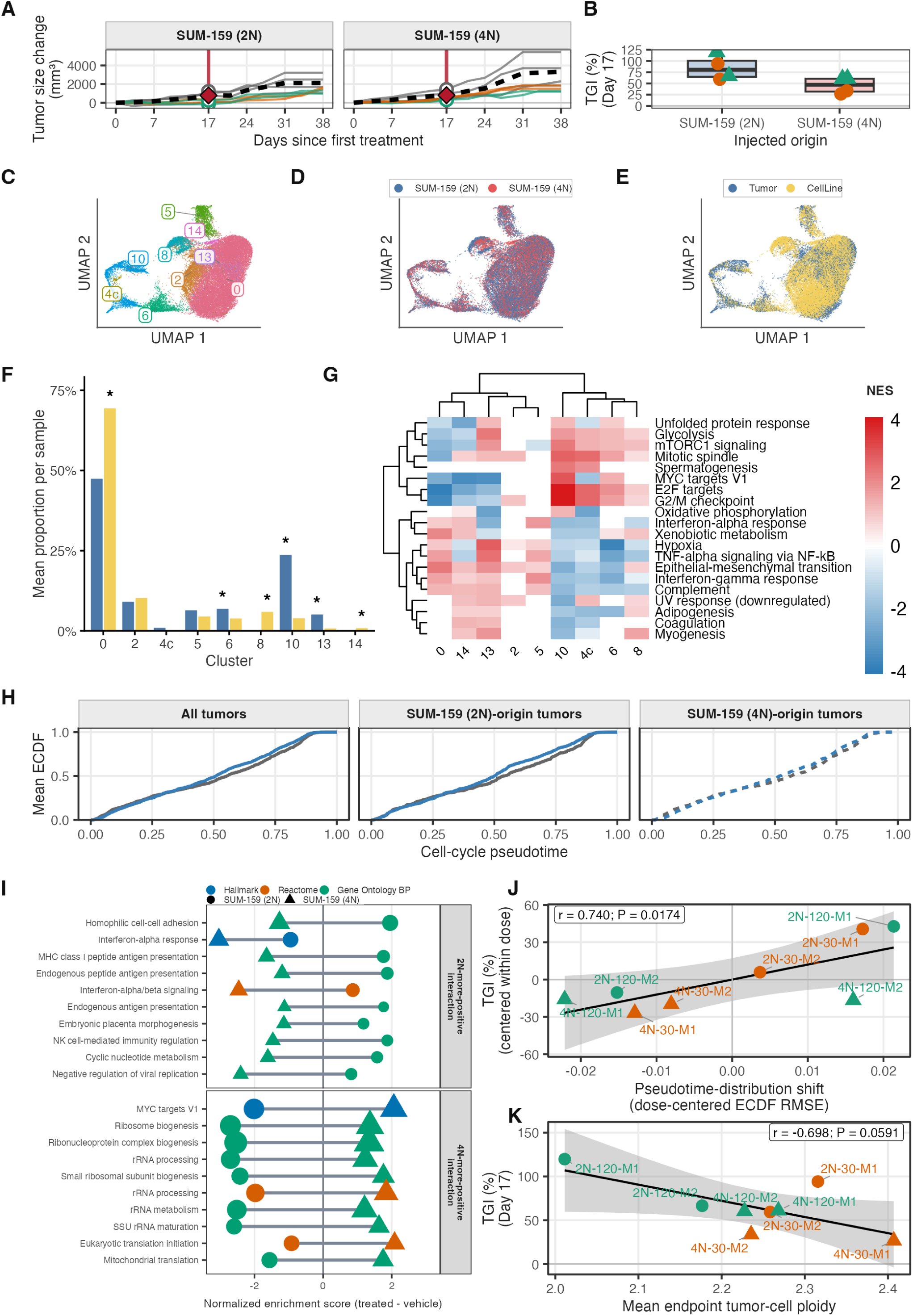
Tumor growth, transcriptional states, and endpoint copy-number architecture in gemcitabine-treated SUM-159 xenograft. **(A)** Change in tumor volume from the first treatment day for tumors initiated with near-diploid (2N) or near-tetraploid (4N) SUM-159 cells and treated with vehicle or gemcitabine at 30 or 120 mg/kg. Thin lines represent individual tumors; dashed black lines show the matched vehicle-group means, and the red vertical line marks Day 17. **(B)** Day-17 tumor growth inhibition (TGI) for gemcitabine-treated tumors grouped by injected ploidy origin. Colors indicate 2N- and 4N-origin tumors, and symbols indicate gemcitabine dose. All tumors are shown. **(C)** Joint uniform manifold approximation and projection (UMAP) of 9,832 tumor cells and 25,681 cultured SUM-159 reference cells, colored by transcriptional cluster. **(D)** The same UMAP colored by injected origin for tumor cells or ploidy of the cultured reference. **(E)** The same UMAP colored by source context. **(F)** Mean within-sample cluster proportions in tumors and cultured references, with each biological sample weighted equally. Asterisks indicate clusters significantly enriched in the context represented by the taller bar after panel-wide multiple-testing correction. **(G)** Cluster-versus-rest Hallmark gene-set enrichment analysis for the nine transcriptional clusters. Red and blue indicate positive and negative normalized enrichment scores (NES), respectively; white tiles denote cluster–pathway combinations without a finite NES. **(H)** Mouse-balanced empirical cumulative distribution functions of cell-cycle-associated pseudotime in vehicle- and gemcitabine-treated tumors. Gray and blue curves denote vehicle and gemcitabine treatment, respectively; solid and dashed curves show the combined and injected-origin-stratified comparisons. The lower strip shows the treated-minus-vehicle density contrast and the marked pseudotime interval enriched in treated tumors. **(I)** Treatment-associated pathway enrichment within the treated-cell-enriched pseudotime interval in 2N- and 4N-origin tumors. The upper panel shows five pathways significant in both origins; lower panels show the most significant nonshared pathways for each origin. Positive and negative NES indicate enrichment in gemcitabine-treated and vehicle tumors, respectively; colors denote gene-set collection and point size reflects FDR. **(J)** Association across treated tumors between Day-17 TGI and displacement of the cell-cycle-associated pseudotime distribution from the matched vehicle reference. Colors indicate dose and symbols indicate injected origin. **(K)** Association across treated tumors between Day-17 TGI and mean endpoint tumor-cell ploidy. Colors and symbols are as in K. Because the 2N- and 4N-origin cohorts were processed in separate NUMBAT analyses, this association is descriptive. TGI, tumor growth inhibition; UMAP, uniform manifold approximation and projection; NES, normalized enrichment score.

We next asked whether the difference in gemcitabine response persisted despite this marked narrowing of the ploidy separation.Tumor growth inhibition (TGI) was calculated on Day 17 from the change in tumor volume after treatment initiation, relative to the mean change of the four vehicle-treated tumors matched by injected origin (Fig. 7A-B). Mean Day-17 TGI was 85.0% in 2N-origin tumors and 45.5% in 4N-origin tumors. The dose-adjusted 4N-minus-2N difference was *−*39.5 percentage points (exact dose-stratified permutation *P* = 0.0556; Fig. 7B). TGI therefore remained lower in 4N-origin tumors, concordant with the direction observed in vitro, even though the realized ploidy separation had narrowed substantially by harvest.

We next examined whether gemcitabine altered tumor-cell composition and transcriptional states within tumors. Joint analysis of 9,832 tumor cells and 25,681 matched cultured SUM-159 reference cells defined a nine-cluster transcriptional landscape (Fig. 7C–E). After calculating cluster proportions within each biological sample and weighting samples equally, tumors showed positive enrichment of clusters 6, 10, and 13, whereas cultured references enriched for clusters 0, 8, and 14 (initial-ploidy-stratified exact permutation, panel-wide Benjamini–Hochberg *q* = 0.037 for each; Fig. 7F, Supplementary Fig. 7A-C). Hallmark pathway analysis nominated clusters 4c, 6, and 10 as cell-cycle associated states with enrichment of complementary E2F-target, G2/M-checkpoint, and mitotic-spindle programs (Fig. 7G, Supplementary Fig. 8A). Gene-set enrichment analysis (GSEA) independently yielded positive normalized enrichment scores (NES) for all three programs in each cluster (Fig. 7G). Clusters 6 and 10 also exhibited high S-phase scores (Supplementary Fig. 9A). We therefore defined clusters 4c, 6 and 10 as an operational cell-cycle-associated subset, comprising 2,881 xenograft-derived cells (Supplementary Fig. 9B), and ordered these cells along a stochastic RNA-velocity-derived pseudotime trajectory rooted in cluster 6.

Gemcitabine treatment produced a reproducible redistribution along this cell-cycle-associated coordinate. Equal-mouse mean empirical cumulative distribution functions (ECDFs) of cell-cycle-associated pseudotime differed between the vehicle and gemcitabine-treated groups (eight tumors per treatment group; RMSE = 0.0378, exact permutation *P* = 0.00513; Fig. 7H). The shift had the same direction within both injected-origin strata, although the smaller comparisons were not independently significant (2N origin: RMSE = 0.0456, *P* = 0.0571; 4N origin: RMSE = 0.0317, *P* = 0.143). In a complementary mouse-balanced density analysis, the treated-minus-vehicle density contrast reached its maximum at pseudotime 0.452. A broader connected region spanning 0.296–0.486 had pointwise permutation support ( *P ≤* 0.05; Supplementary Fig. 9C), further referred to as *treated-cell-enriched pseudotime interval*. To characterize treatment-associated transcriptional programs within this interval, we compared gemcitabine-treated and vehicle-treated tumors using mouse-level pseudobulk models fit separately for 2N- and 4N-origin tumors and used the resulting gene rankings for pathway enrichment analysis (Fig. 7I). Within the selected pseudotime interval, SUM-159 (2N)-origined tumors showed stronger suppression of ribosome production and RNA/protein biosynthetic capacity, whereas the SUM-159(4N)-origined tumors were characterized predominantly by reduced interferon signaling, and antigen processing and presentation programs. Together, gemcitabine was associated with substantially origin-dependent transcriptional programs within the *treated-cell-enriched pseudotime interval*, with 2N-derived tumors showing more extensive reduction of anabolic and proliferative programs in response to gemcitabine compared to their 4N counterparts. The extent of this transcriptional redistribution was related to tumor response. Across the eight treated tumors, pseudotime-distribution displacement from the equal-mouse vehicle reference matched by injected origin was positively associated with Day-17 TGI after both quantities were centered within dose (*r* = 0.740, exact within-dose permutation *P* = 0.0174; Fig. 7J). Among these same tumors, Day-17 TGI showed a sizeable inverse association with mean endpoint tumor-cell ploidy (Pearson *r* = *−*0.698, exact unrestricted permutation *P* = 0.0591; Fig. 7K). This association is consistent with stronger inhibition at lower realized ploidy, but the eight-mouse sample and limited endpoint-ploidy variation constrain precision. Sensitivity analyses at later endpoints preserved these directions but attenuated their magnitudes (Day 24 TGI: *P* = 0.167; Day 31 TGI: *P* = 0.306; Supplementary Fig. 10A,B,D). Endpoint-ploidy correlations also remained inverse but were weaker on Day 24 (*r* = *−*0.387, *P* = 0.3469) and Day 31 (*r* = *−*0.297, *P* = 0.4857; Supplementary Fig. 10C,E).

Taken together, mean early TGI remained substantially lower in 4N-origin tumors despite pronounced narrowing of the ploidy difference. Gemcitabine also produced a significant redistribution along cell-cycle-associated pseudotime, and the magnitude of this redistribution tracked early tumor-growth inhibition within dose. The *treated-cell-enriched pseudotime interval* defined a replication- and biosynthesis-attenuated state along the cell cycle trajectory. Endpoint convergence provides biological context for the limited endpoint-ploidy variation and precision of the in vivo association. Within that compressed range, lower endpoint ploidy was associated with greater gemcitabine response in the same direction observed in the controlled in vitro experiments.

## Discussion

Our study identifies ploidy as a determinant of both the magnitude and kinetics of gemcitabine response. Across independent pharmacogenomic datasets, gemcitabine repeatedly emerged as a low-ploidy-selective cytotoxic agent, and this association was recapitulated across matched isogenic systems from multiple lineages. Our mechanistic analysis of near-diploid and near-tetraploid isogenic cell line pair further revealed that high-ploidy resistance was not explained by a failure to generate the active metabolite dFdCTP. Instead, 4N cells accumulated more dFdCTP while exhibiting weaker checkpoint activation, attenuated nucleotide-metabolic remodeling, reduced fitted cytostatic and cytotoxic potency, and a substantially longer delay before cell death. These findings support the SUM-159 live/dead model in which high ploidy alters the pharmacodynamic conversion of intracellular gemcitabine exposure to weaker replication stress, cytostatic transition and cytotoxic potency together with a substantially longer delay from effective intracellular dFdCTP exposure to observed cell death. Thus, the current data indicate a parsimonious interpretation in which ploidy reshapes gemcitabine response through altered effective drug-action mapping and delayed execution of cell death.

The dissociation between dFdCTP accumulation and cytotoxic response is a central feature of our study. Canonical models of gemcitabine resistance frequently invoke reduced transport, impaired phosphorylation, enhanced inactivation or diminished intracellular retention. Our results indicate that active-metabolite abundance alone is insufficient to predict drug response across ploidy states. High-ploidy cells remained comparatively resistant despite accumulating substantial intracellular dFdCTP, suggesting that equivalent or even greater total metabolite exposure does not necessarily impose equivalent functional pressure on DNA synthesis. Ploidy may, therefore, influence gemcitabine response downstream of metabolic activation, through differences in the scale of nucleotide demand, the effective competition between dFdCTP and endogenous dCTP, the number and behavior of replication forks or the capacity to sustain DNA synthesis under nucleotide stress.

The checkpoint signaling provides a mechanistic anchor for the model-inferred ploidy-dependent potency shift. Gemcitabine induced stronger phosphorylation of CHK1, and to a lesser extent CHK2, at lower administered doses in 2N cells compared to 4N cells. Together with the intracellular PK/PD and live-cell measurements, this result indicates that lower-ploidy cells translate gemcitabine exposure into detectable replication-stress signaling more efficiently. However, checkpoint activation is not itself a direct measure of lethal damage because it can reflect both the magnitude of replication stress and the protective response mounted against it. Therefore, reduced checkpoint signal in 4N cells could arise from weaker initial nucleotide stress, delayed pathway engagement, enhanced replication-fork stabilization or altered tolerance of lesions that persist into mitosis.Future time-resolved and cell-cycle-resolved measurements will, thus, be needed to determine whether 4N cells experience less initial replication stress, delayed checkpoint activation, enhanced replication fork protection or greater tolerance of damage carried into mitosis.

The metabolomic data suggest that differential nucleotide homeostasis contributes to this phenotype. Gemcitabine produced broader metabolic remodeling in 2N cells, including stronger changes in pyrimidine- and purine-associated metabolites and a larger reduction in dCMP abundance whereas 4N cells showed a comparatively attenauted response. These findings suggest that gemcitabine sensitivity depends on the balance between demand, competing cytidine pools and nucleotide-buffering capacity. Proliferative 4N cells that have successfully adapted to genome doubling must support replication of an expanded DNA template and may consequently acquire increased nucleotide-production or salvage capacity to help buffer gemcitabine-induced cytidine-pathway perturbation. Therefore, such adaptations may incidentally reduce the effective replication-stress burden imposed by gemcitabine. Here, high-ploidy cells could be more resistant if they maintain a lower effective dFdCTP-to-dCTP pressure, either by accumulating larger competing cytidine pools, sustaining nucleotide replenishment, or reducing the functional impact of gemcitabine metabolites on DNA synthesis. This interpretation is also important for evaluating intracellular dFdCTP measurements: because 4N cells are larger, greater total dFdCTP accumulation in 4N cells (Fig. 4B), need not imply greater effective drug pressure per unit DNA, cell volume, replication fork, or competing cytidine nucleotide pool. However, because steady-state dCMP abundance does not directly measure dNTP pools or nucleotide flux, future isotope-tracing and targeted dNTP/dFdCTP measurements will be required to distinguish increased de novo synthesis from altered salvage, catabolism, nucleotide utilization or scaling effects related to cell size and DNA content.

A second, non-mutually exclusive interpretation is that higher-ploidy cells may also be better able to tolerate the chromosomal consequences of replication-stress-induced missegregation. Under-replicated DNA becomes most dangerous when cells enter mitosis before replication is complete, because unresolved loci can generate ultrafine anaphase bridges, chromatin bridges, acentric fragments, micronuclei and post-mitotic genome loss [11–13]. In this framework, high ploidy is not expected to be uniquely compatible with upstream pharmacologic resistance mechanisms, which primarily reduce effective intracellular drug burden. Instead, high ploidy is most compatible with downstream resistance mechanisms that require survival after imperfect genome duplication, including tolerance of under-replicated DNA entering mitosis, bridge or micronucleus resolution, senescence or PGCC-like persistence, and viable exit from abnormal mitoses [1, 10, 39, 40]. Thus, our results suggest that gemcitabine resistance can arise not only by reducing the initial replication lesion, but also by increasing the probability that cells survive the chromosomal damage produced when replication stress is carried into mitosis.

Many anticancer drugs promote chromosome missegregation either directly, by disrupting mitosis, or indirectly, by generating replication and DNA-damage lesions that persist into mitosis. A ploidy-dependent survival framework predicts that near-diploid cells should be particularly vulnerable to CIN-inducing therapies, because missegregation events, especially chromosome-loss events, impose larger immediate fitness costs in cells with less DNA copy number redundancy. Consistent with this prediction, previous pharmacogenomic analyses identified multiple mitotic spindle poisons, including paclitaxel, vinblastine, and vinorelbine, among the strongest ploidy-associated agents in breast cancers [16]. Surprisingly, gemcitabine showed the strongest association with ploidy in this analysis. Although gemcitabine is classically viewed as an S-phase antimetabolite, an independent human artificial chromosome loss assay showed that it is also a potent inducer of chromosome loss, ranking among the highest CIN-inducing drugs tested and increasing chromosome loss by approximately 35–45-fold [14]. Independent drug-sensitivity studies have also identified gemcitabine as a low-ploidy-selective compound [18]. These observations nominate gemcitabine as an unexpected but compelling candidate for testing whether ploidy modifies chemotherapy response.

Our mathematical model formalizes this distinction between intracellular exposure and downstream effect. A model driven solely by the measured dFdCTP profiles was insufficient to reproduce the observed responses across doses and ploidy states. Incorporating a nonlinear dose transformation, ploidy-specific potency parameters and a distributed delay to death substantially improved the representation of the live and dead cell trajectories. The fitted model attributed the enhanced sensitivity of 2N cells to both stronger immediate cytostasis and greater delayed cytotoxicity, while estimating a more than twofold longer interval between effective drug action and observed death in 4N cells. These parameters provide a quantitative summary of otherwise unresolved biological steps separating metabolite exposure from phenotypic outcome and identify the exposure-to-effect relationship—not drug accumulation alone—as the principal point of divergence between ploidy states. Moreover, this mathematical approach indicated that high-ploidy cells may translate intracellular gemcitabine exposure into a more attenuated metabolic response. Guided by this prediction of the model, metabolomic profiling revealed broader disruption of nucleotide-associated pathways in 2N cells than in 4N cells, providing a plausible metabolic basis for the stronger replication-stress response observed in the low-ploidy state.

Altogether, our findings establish an experimental and mathematical framework in which ploidy functions as a quantitative cell-state variable governing gemcitabine response. High-ploidy cells do not simply accumulate less active drug; rather, they appear to buffer or delay the processes that connect dFdCTP exposure to nucleotide disruption, checkpoint activation, and cell death. These findings have implications beyond gemcitabine. For therapies whose activity depends on DNA synthesis, genome copy number may determine both the effective damage caused by treatment and the probability that damaged cells survive long enough to recover, persist, or re-enter the cell cycle. Incorporating ploidy into pharmacodynamic analyses may therefore reveal therapeutic dependencies that remain obscured when drug response is interpreted solely from extracellular dose or intracellular drug abundance.

## 1 Methods

### 1.1 Cell lines

TNBC lines SUM-159 and MDA-MB-231 were a generous gift of Andriy Marusyk, and gastric cancer lines SNU-668 and HGC-27 were obtained from the ATCC. Cell-line identities were confirmed by short tandem repeat analysis of cellular DNA, and all cultures were routinely tested for Mycoplasma contamination. TNBC lines were cultured in McCoy medium (GIBCO, 16600), and gastric cancer lines were cultured in RPMI-1640 medium (GIBCO, 11875) supplemented with 10% FBS (v/v; Gemini Bio, 900-108-500).

Isogenic low- and high-ploidy cancer-cell pairs were generated by long-term density selection in SNU-668 [41] and by cell fusion [42] and long term hypoxia (unpublished) in SUM-159. Additional matched low- and high-ploidy lineage pairs were included as indicated. For each isogenic pair, DNA-content differences were confirmed by flow cytometry and maintained for at least five passages before drug exposure.

### 1.2 Reagents

Gemcitabine (ChemieTek, CT-GEM) was purchased in powder form and stored at 10 or 20 mmol/L stock concentrations in DMSO at *−*20*^◦^*C. Phospho-CHK1 (Ser317; Cell Signaling Technology, rabbit mAb, 12302, 1:1,000 dilution), Phospho-CHK2 (Thr68; Cell Signaling Technology, rabbit mAb, 2661, 1:1,000 dilution), *β*-actin (MP Biomedicals, mouse mAb, 69100, 1:10,000 dilution) antibodies were used for immunoblotting assays.

### 1.3 Pharmacogenomic analysis of ploidy-associated drug classes

We integrated Genomics of Drug Sensitivity in Cancer (GDSC) drug-response data [33] with cell-line ploidy estimates from Cell Passports [34]. For each cancer type and drug, duplicate drug–cell-line records were resolved by retaining the measurement with the lowest finite root mean squared error (RMSE). We then calculated the Pearson correlation between cell-line ploidy and the GDSC drug-response Z-score across matched cell lines. Drugs were assigned to 19 primary anticancer classes derived from Pubchem and Drugbank [43]; then grouped into 11 broader displayed categories. We then tested whether each drug category was enriched near the top of the signed ranked list using a one-sided rank-enrichment statistic analogous to a Kolmogorov–Smirnov running-sum statistic. Enrichment significance was estimated with 20,000 random permutations of drug-category labels. Exact zero permutation *p*-values were replaced by the permutation floor 1/(20, 000 + 1), and Benjamini–Hochberg false discovery rate (BH-FDR) correction was applied across the displayed drug-category, cancer-context and enrichment-direction tests. Figure 1 reports enrichment at BH-FDR *q* ≤ 0.05

#### 1.3.1 CCLE ploidy-associated drug sensitivity analysis

To locally reproduce the breast cancer CCLE ploidy–drug-response analysis, we matched breast cancer cell-line ploidy estimates to the legacy Breast Cancer Drug Sensitivity drug-response tables used by the original local analysis. These tables report drug-response *Z*-scores [16, 17, 44]. Replicate or repeated drug entries within a cell line were averaged, and drugs were retained if they were observed in at least 80% of the maximum drug coverage across matched cell lines. We then computed the Pearson correlation between cell-line ploidy and drug-response *Z*-score across matched cell lines. Lower *Z*-score values were treated as indicating greater drug sensitivity; therefore, positive correlations indicate that higher-ploidy lines tend to have higher *Z*-scores, consistent with relatively greater sensitivity in lower-ploidy lines. For visualization, retained drugs with *|r| ≥* 0.2 were plotted.

### 1.4 Microplate drug-sensitivity assays

SUM-159, MDA-MB-231, SNU-668 and HGC-27 cells (3,000 per well) were seeded in 96-well plates and incubated at 37*^◦^*C for 24 hours. Treatments with vehicle (DMSO) or serial dilutions of the indicated compounds in 11 concentrations from 1 to 10,000 nmol/L were performed in triplicate (final DMSO concentration = 0.34% in all wells). After 96 hours, media were removed and cell viability was measured with a CellTiter-Glo assay (Promega, G7573).

For the Figure 3H dose-response/AUC analysis, sample-level responses were normalized to the corresponding untreated control mean. Four-parameter log-logistic dose-response curves were fit with `drc::LL.4`, and normalized AUC was calculated over a common log_10_ dose interval from 0.005 to 0.9 *µ*M. Mean sample ploidy was taken from CLONEID genome-perspective profiles when available; documented fallback ploidy values were used for legacy samples without a matched CLONEID profile.

### 1.5 Crystal violet staining assay

SUM-159 cells (1 *×* 10^5^ per well) were seeded in 12-well plates and incubated at 37*^◦^*C for 24 hours. Treatments with vehicle (DMSO) or serial dilutions of the indicated compounds in 10 indicated concentrations were performed (final DMSO concentration = 0.34% in all wells). After seven days, media were removed and cell growth was assessed with a crystal violet staining assay solution [6% glutaraldehyde, 0.5% crystal violet (w/v) in water].

### 1.6 Light microscope and immunofluorescence assay

Cells in transparent bottom plates were used to monitor cell morphology and nuclear mCherry expression. Microscopic observations were performed on a Nikon or EVOS FL Cell Imaging System. Mean nuclear area of cells after the treatments with the indicated compounds were calculated with the ImageJ software and graphed. RGB images of cells were converted to CMYK, and visualized with monochromic magenta (red and blue) or cyan (green and blue) filters to reduce the endogenous autofluorescence and acquire a detailed view of the cellular structures of interest such as nucleus and centrosome, if indicated.

### 1.7 Delay-aware pharmacodynamic (PD) model of gemcitabine response

Let *p ∈ {*2*N,* 4*N }* denote ploidy state and let *C_p_*(*t, d*) denote the fitted intracellular dFdCTP-like exposure signal for dose *d* (Supplementary Methods 2.1.1). The delayed drug-effect signal is represented by the terminal compartment *Z_n,p_*(*t, d*) of an *n*-stage transit chain driven by *C̄p*(*t, d*):

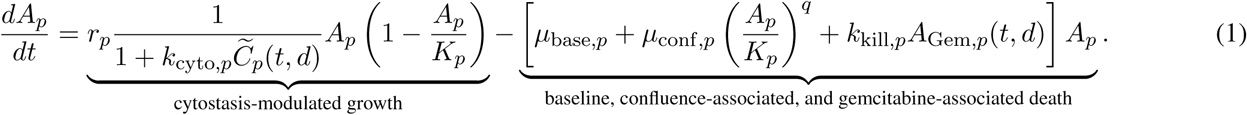

Here, *A_p_*(*t*) denotes live-cell abundance, *r_c_* is baseline growth rate, *K_p_* is carrying capacity, *k*_cyto*,p*_ controls apparent cytostatic potency, *µ*_base*,p*_ and *µ*_conf*,p*_ capture baseline and confluence-associated death, and *k*_kill*,p*_ controls apparent cytotoxic potency. The transit-chain construction captures distributed delay from intracellular drug activity to observed death [45, 46].

### 1.8 Incucyte-based cell viability assay

Cell viability assays were carried out using the IncuCyte live-cell imaging system (Sartorius). Briefly, 2N and 4N SUM-159-NLS-mCherry cells were plated in four replicates in 96-well plates (Falcon, Corning, New York, USA) at a density of 2000 cells per well and treated with a 10-point 2X serial dilutions of gemcitabine within the concentration range of 0 and 800 nM. Sytox Deep Red (S11381, ThermoFisher Scientific) nucleic acid stain was also added to monitor dead cells. Plates were then incubated in the IncuCyte CO2 incubator for 10 days while four separate images were captured from each well every 2 hours using 10× objective in red, infrared and phase channels. Red object count (ROC) per well was calculated as a measure of viable cells and near-infrared object counts (NOC) per well was calculated as a measure of dead cells when the treatment was performed.

### 1.9 Quantification of gemcitabine metabolites by LC–MS/MS

Matched near-diploid (2N) and near-tetraploid (4N) SUM-159 breast cancer cells were treated with gemcitabine before submission to the PK/PD Core. Cells were exposed to either 0.1 *µ*M gemcitabine and collected at 0, 0.25, 0.5, 1, 2, and 24 h, or 1.0 *µ*M gemcitabine and collected at 0, 1, 2, 24, and 48 h. Harvested cell pellets containing approximately 1 *×* 10^6^ cells per sample were submitted for targeted quantification of gemcitabine (dFdC) and its metabolites dFdU and dFdCTP.

Cell pellets were resuspended in 1 mL of quenching solution consisting of acetonitrile (50:50, v/v) containing 100 *µ*g*/*mL tetrahydrouridine (THU). A 100 *µ*L aliquot of each sample was supplemented with stable-isotope-labeled internal standards (^13^C^15^N-gemcitabine and ^13^C^15^N-dFdU), followed by protein precipitation and analyte extraction with 300 *µ*L of pre-chilled acetonitrile. Samples were vortex-mixed and centrifuged at 12,500 rpm for 5 min at 4 *^◦^*C using an Eppendorf 5417R refrigerated microcentrifuge (Eppendorf, Hamburg, Germany). The resulting supernatant was transferred to a 96-well plate and evaporated to dryness under nitrogen at 45 *^◦^*C using a Porvair Ultravap nitrogen evaporator (Porvair Sciences, Wales, UK). Dried extracts were reconstituted in 100 *µ*L of 10 mM ammonium acetate containing 0.1% ammonium hydroxide.

Calibration curves for dFdC, dFdU, and dFdCTP were prepared in cell matrix by adding working standard solutions of known concentration to cell homogenate. The calibration range was 1.56–400 ng/mL. Quality-control samples were prepared at 4.7, 18.7, and 75 ng/mL. Calibration standards and quality-control samples were processed using the same sample-preparation procedure as the experimental samples and subsequently analyzed by LC–MS/MS.

A 10 *µ*L aliquot was injected onto a Vanquish UHPLC system coupled to a TSQ Altis Plus triple-quadrupole mass spectrometer (Thermo Scientific, San Jose, CA). Chromatographic separation was performed using an ACQUITY BEH C18 column (2.1 *×* 30 mm, 1.7 *µ*m). Mobile phase A consisted of 10 mM ammonium acetate containing 0.1% ammonium hydroxide, and mobile phase B consisted of methanol containing 0.1% ammonium hydroxide. Analytes were separated using a 4-min gradient at a flow rate of 0.4 mL*/*min.

Gemcitabine and dFdCTP were detected in positive-ion mode using selected-reaction-monitoring transitions *m/z* 264.085 *→* 112.060 and 504.010 *→* 326.062, respectively. dFdU was detected in negative-ion mode using the transition *m/z* 263.045 *→* 219.982. Stable-isotope-labeled gemcitabine and dFdU were monitored at *m/z* 267.120 *→* 115.047 and 266.070 *→* 220.946, respectively. Quantification was performed using TraceFinder software, version 5.2 (Thermo Scientific).

Calibration curves were generated by linear regression with 1*/x*^2^ weighting. Calibration standards and quality-control samples were considered acceptable when their back-calculated concentrations were within *±*15% of their nominal concentrations, except at the lower limit of quantification (LLOQ), for which deviations of up to *±*20% were permitted. “N/F” indicates that no analyte peak was observed; “BDL” indicates that the analyte concentration was below the established detection limit of 0.4 ng/mL; and “BQL” indicates that the analyte was detected but was below the LLOQ of 1.56 ng/mL. The analytical approach was informed by previously described methods for quantifying intracellular gemcitabine triphosphate in cancer cells [**?**].

The source document leaves the second component of the “acetonitrile (50:50, v/v)” quenching solution unspecified. That composition should be confirmed before publication.

### 1.10 PK/PD measurement of intracellular dFdCTP

To construct the intracellular drug-action input for the live/dead model, we used PK/PD measurements of gemcitabine metabolites collected for the matched 2N and 4N cell states. The live/dead model was driven by the intracellular active metabolite dFdCTP, rather than by parent gemcitabine concentration. For each ploidy, PK/PD sheets corresponding to the main-dose and low-initial-dose gemcitabine conditions were used to define calibrated dFdCTP time courses. The main 2N and 4N PK/PD sheets were treated as 1.0 *µ*M gemcitabine reference-dose profiles, whereas the low-initial-gemcitabine 2N and 4N sheets were treated as 0.1 *µ*M reference-dose profiles.

PK/PD dFdCTP measurements were reported in ng/mL and converted to *µ*M-equivalent concentrations using the dFdCTP molecular weight,

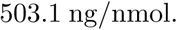

Because

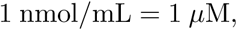

the conversion was

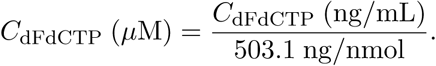

Measurements reflect analyte abundance per approximately (10^6^) cells extracted using a fixed volume, without further cellular normalization, rather than direct intracellular molar concentrations. N/F, BDL, and BQL results were reported categorically without numerical concentration assignment. The resulting dFdCTP profiles were baseline-subtracted using the first measured time point so that the model input represented treatment-induced intracellular dFdCTP above the initial measured level.

### 1.11 Image analysis

#### 1.11.1 Cell segmentation and classification with HALO

Cell segmentation was achieved by training a single-cell segmentation neural network in Halo-AI platform [47]. At the beginning, a training dataset was collected by selecting a subsample of images from the data. Two main selection criteria were considered during the image sampling. First, the training set must have representative images from both 2N and 4N cells. Second, multiple images were sampled from different time points for each cell type 2N and 4N to ensure the capture of cell change over time during the training process. After the selection of train set, nucleus of cells was labeled using Halo built-in labeling tools. Finally, the training process was performed until all cells are accurately segmented. Another object classifier was trained on the top of the cell segmentation classifier results to classify each segmented cell to either dead, alive or transitional cell. At the beginning, cells are labeled either dead or alive by tracking the cells over the time and by observing changes in their nuclei. The object classifier was trained iteratively until the cell were classified correctly.

### 1.12 Flow cytometry analysis

For the DNA content analyses, 10^6^ cells were resuspended in 700 µl of PBS, and then 700ul of 8% formaldehyde was added dropwise. The cells were incubated for 30 minutes, then washed in PBS twice and resuspended in 300µl of PBS with 0.2% Triton X-100 (ThermoFisher) and 5 uM Vybrant DyeCycle Violet (ThermoFisher). The flow cytometry analysis was performed using a FACSCanto II instrument (BD Biosciences), and the data were analyzed using FlowJo 10.10.1 software. The average number of collected events was 30,000. For each of the tested samples, flow cytometry analyses were performed over two or more distinct experiments.

### 1.13 Western blotting

Cells were seeded in 6-well plates at 2 × 10^5^ cells per well and treated with the indicated compounds at the indicated concentrations. 24 hours after treatment, culture medium was aspirated and cells were washed once with cold PBS. Cells were lysed in ice-cold RIPA buffer containing 50 mmol/L Tris-HCl pH 8.0, 150 mmol/L NaCl, 1% NP-40, 0.1% SDS, and 0.5% sodium deoxycholate, supplemented with protease and phosphatase inhibitor cocktails. Lysates were clarified by centrifugation at 14,000 × g for 10–15 min at 4°C, and protein concentrations were determined using the DC Protein Assay (Bio-Rad, 500-0111).

Equal amounts of protein were resolved by SDS-PAGE and transferred to low-fluorescence nitrocellulose membranes. Membranes were blocked in LI-COR Intercept Blocking Buffer for 1 h at room temperature and incubated overnight at 4°C with primary antibodies against phospho-CHK1, phospho-CHK2, and β-actin as an internal loading control. After washing with PBS-T, membranes were incubated for 1 h at room temperature with species-appropriate IRDye 680RD- or IRDye 800CW-conjugated secondary antibodies protected from light. Membranes were washed again with PBS-T followed by PBS and imaged using a LI-COR Odyssey infrared imaging system.

### 1.14 Untargeted Metabolomics

Ultra high-performance liquid chromatography-high resolution mass spectrometry (UHPLC-HRMS) was performed using a Vanquish UHPLC interfaced with a Q Exactive HF quadrupole-orbital ion trap mass spectrometer (Thermo, San Jose, CA). Chromatographic separation was performed using an Atlantis Premier BEH Z-HILIC VanGuard FIT column (2.1 mm ID x 150 mm length, 2.5 µm particle size, Water, Milford, MA). Mobile phase A was aqueous 10 mM ammonium carbonate and 0.05% ammonium hydroxide, and mobile phase B was 100% acetonitrile. The gradient program included the following steps: start at 80% B, a linear gradient from 80 to 20% B over 13 minutes, stay at 20% B for 2 minutes, return to 80% B in 0.1 minutes, and re-equilibration for 4.9 minutes for a total run time of 20 minutes. The flow rate was set to 0.250 mL/min. The autosampler was cooled to 5 °C and the column temperature was set to 30 °C. Sample injection volume was 2 µL for both positive ion mode and negative ion mode electrospray ionization. Full MS was performed in positive and negative mode separately detecting ions from m/z 65 to m/z 900. In addition, data-dependent acquisition was used for tandem mass spectrometry (MS/MS) of analytes in the pooled samples to enable verification of selected metabolites to confirm assignments. MZmine software, version 2.53, was used to identify and quantify metabolites by matching by m/z and RT to an in-house library. A batch file was used to automate the following modules: centroid mass detection, ADAP chromatogram builder (with 5 scans minimum group size and group intensity threshold and minimum highest intensity set to 1.0E4), smoothing (set to 5), deconvolution by local minimum search (with Chromatographic threshold 95%, 0.05 minutes search minimum in RT range, 10% minimum relative height: 10%, 1.0E4 minimum absolute height, minimum peak top/edge set to 1, and 0.05-5 minutes peak duration), isotopic peak grouping (with 10 ppm m/z tolerance, 0.25 minute Retention time tolerance, and maximum charge state 2), peak alignment (using m/z and RT tolerances from the previous step with 75% weighting for m/z and 25% weighting for RT), peak finder (with 10% intensity tolerance and m/z and RT tolerances set as in the previous steps), gap filling, duplicate peak filtering (in new average mode with m/z and RT tolerances as above), custom database search (using an in-house library with m/z tolerance 10 ppm and RT tolerance 0.3 minutes), adduct and complexes search, peak list row filtering, and peak list export. Peak height values were exported (csv files) for further analysis by BBSR.

### 1.15 Cytogenetics and G-banded karyotyping

To evaluate chromosomal stability and structural integrity, G-banded karyotyping was performed using the commercial service provided by Labcorp Specialty Testing Group (Burlington, NC, USA). Cultured cells were harvested during log-phase growth and treated with colcemid to arrest dividing cells in mitosis. Mitotic cells were subsequently subjected to hypotonic treatment, fixed in a methanol and acetic acid solution (3:1 v/v), and dropped onto clean glass slides to yield optimal metaphase spreads. Chromosomes were treated via enzymatic digestion using trypsin and stained with Giemsa (GTW/GTG banding) to produce distinct, reproducible light and dark horizontal banding patterns. Cytogenetic analysis was conducted by clinically certified cytogenetic technologists, and a minimum of 20 metaphase spreads were microscopically reviewed per sample at a standard resolution of 400–450 bands per haploid set. Final data evaluation and structural interpretation were validated by a certified laboratory director.

### 1.16 Mouse xenograft studies

Cells derived from the isogenic human TNBC lines SUM-159 (2N) and SUM-159 (4N) were trypsinized and collected in the growth media supplemented with 10% fetal bovine serum (v/v). The cells were centrifuged at 150g and the media was aspirated. After washing twice with sterilized PBS, cells were resuspended and diluted in PBS to have 1 × 10^6^ cells per 50 µl concentration. Single-cell suspensions (50 µl) were injected into the mammary fat pads of 8–10-week-old, female NOD/SCID mice. Mice xenografted with SUM-159 cells were kept in a sterile vivarium maintained by the Animal Resource Center at Moffitt Cancer Center. Tumor volumes were measured twice a week with caliper measurements and calculated by the formula: tumour volume=(length × *width*^2^)/2. Gemcitabine treatment was initiated when tumours reached a mean volume of 100–150*mm*^3^. Mice were assigned to treatment groups randomly after sorting according to the tumor volume. Gemcitabine (Pfizer Inc., 0409-0182-01) was formulated with saline (Alpha Teknova Inc., S5825). For each dose of gemcitabine, five mice xenographed with each cell line were used. Mice were injected intraperitoneally with 0, 30, 60 and 120 mg*kg^−^*^1^ gemcitabine twice weekly. The experiments were terminated when the largest tumours reached 1500 *mm*^3^. Animal studies were carried out under a protocol approved by the Institutional Animal Care and Use Committee of UT Southwestern that conforms to the requirements of Animal Act PL99-158 (as amended) and guidelines stated in the ‘Guide for the Care and Use of Laboratory Animals’.

### 1.17 Viable tumor dissociation

Single cell isolates from fresh collected tumors were obtained using the Miltenyi Human Tumor Dissociation kit (Cat. #130-095-929) and the gentleMACS Dissociator (Cat.#130-095-235) according to manufacturer’s instruction. Tissues were grossed to remove fatty, fibrous and necrotic areas. The tissue sample (up to 200 mg) is then minced in a glass petri dish using a sterile scalpel and transferred to a gentleMACS C tube (Cat. #130-093-237). A prepared proprietary enzyme mixture of kit supplied enzyme H, R, and A were then added to the minced tissue sample and the tissue-enzyme suspension was dissociated using the gentleMACS Dissociator for 3 cycles of Program h_tumor_02 followed by a 30-minute incubation at 37*^◦^*C under continuous rotation. The dissociated cell suspension was then passed through a MACS 70um SmartStrainer (Cat. #130-110-916), washed with RPMI medium, resuspended in CS10 cryopreservation medium (CryoStor, Cat. #C3124, Sigma), and used immediately or stored at -80*^◦^*C until use.

### 1.18 In-vivo derived scRNA-seq analysis

Single-cell RNA sequencing was performed on 16 primary tumors selected to include all vehicle-treated tumors and representative samples from treatment groups. The two parental cell lines, SUM-159 (2N) and SUM-159 (4N), were also sequenced as references. Raw sequencing data was processed using the 10x Genomics Cell Ranger pipeline to generate feature-barcode matrices. As the experiment was conducted using an orthotropic mouse model, raw data was aligned to a combined human (GRCh38) and mouse (mm10) reference genome. Each cell was then classified as either human (tumor) or mouse (tumor microenvironment) based on the species to which the majority of its UMIs aligned. Only cells unambiguously assigned to one species were retained.

For each sample, further quality control was performed independently. Low-quality cells were identified and removed based on standard metrics, including the number of unique molecular identifiers (UMIs), the number of detected genes, and the percentage of mitochondrial gene expression. Specifically, cells with an unusually low or high number of detected genes, or with a mitochondrial gene content exceeding 10%, were excluded from downstream analysis to remove empty droplets, cellular debris, and stressed or dying cells. Following filtering, the gene expression data for each sample was normalized using the `SCTransform` method in the Seurat R package (v4), which stabilizes variance and identifies high-variance features.

#### 1.18.1 Copy number analysis

##### Numbat

Single-cell copy number profiles were inferred using Numbat. For the diploid setting, 2N tumor cells was analyzed together with a 10% spike-in of 2N cell line cells, whereas for the tetraploid setting, 4N cells were analyzed with a 10% spike-in of 4N cell line cells. Reads mapped to the human reference genome (GRCh38) were retrieved from 10x Genomics BAM files and used to compute phased allele counts at single-cell resolution, using haplotype information from the 1000 Genomes Project as the reference panel. Copy number states were estimated by jointly modeling allele-specific counts and gene expression matrices within the Numbat framework, using the following parameters: *t*=1e-5, *initial k*=3, *min genes*=10, *max entropy*=0.5. Chromosome 2 was specified as the diploid reference for 2N cells, and Chromosome 13 was specified as the diploid reference for 4N cells. A diploid reference derived from the Human Cell Atlas was applied in both settings.

##### Bayesian Shrinkage Adjustment of Segmental Copy Number by Expression

For each genomic segment *i* with an a priori copy number estimate 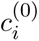, we refined copy number values by integrating expression data derived from matched RNA profiles. Gene-level normalized expression values were first smoothed using a 400-gene rolling mean across genomic coordinates. Each gene was then assigned to the segment(s) it overlapped, and the mean rolling expression per segment (*e_i_*) was computed. To relate expression and copy number on a genome-wide scale, we fit a robust linear model

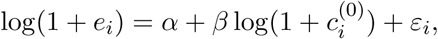

using Huber weighting to down-weight outlier segments (`MASS::rlm` with *ψ* = psi.huber*, k* = 1.5). For each segment, the expression-implied copy number was obtained by inverting the fitted relationship:

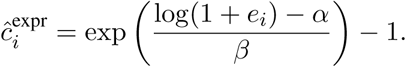

Uncertainty of this estimate was approximated using the delta method,

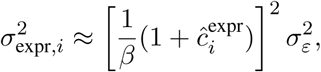

where 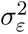 is the residual variance of the regression. An empirical prior variance 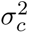 was computed from the deviation between prior and expression-implied copy numbers, 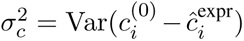. For each segment, the maximum a posteriori (MAP) adjusted copy number was then obtained by Bayesian shrinkage toward the prior:

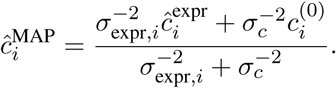

This formulation balances expression-derived evidence against the genomic prior according to their respective uncertainties. To prevent biologically implausible values, adjusted copy numbers were truncated to the range [1, 6]. The procedure returns both the fitted regression parameters 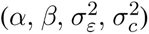 and a per-segment table containing prior, expression-implied, and MAP-adjusted copy numbers for downstream analysis.

## Data and code availability

Processed single-cell RNA-sequencing data are available through Zenodo (DOI: 10.5281/zenodo.21463392) including sample-level filtered gene-expression matrices, spliced and unspliced RNA counts, and the integrated Seurat object used for downstream analyses. Analysis-ready datasets and code are publicly available in the Gemcitabine-model GitHub repository (https://github.com/noemiandor/Gemcitabine-model). These include the public pharmacogenomic inputs and curated drug classifications used in Figures 1–2; gemcitabine dose-response and ploidy measurements for Figure 3; live-cell imaging count tables and intracellular drug measurements for Figures 4–5; metabolite-abundance measurements for Figure 6; and tumor-growth measurements, sample metadata, single-cell analysis tables, copy-number estimates, and endpoint flow-cytometry data supporting Figure 7 and the supplementary figures. A figure-by-figure dataset guide in Data/README.md identifies the corresponding files, their provenance, and the workflows used to generate the agreedfigures.

## Acknowledgements

This work has been supported in part by the Cancer Pharmacokinetics and Pharmacodynamics, Flow Cytometry, Molecular Genomics, Tissue, and the Proteomics and Metabolomics Core Facilities at the H. Lee Moffitt Cancer Center & Research Institute, an NCI designated Comprehensive Cancer Center (P30-CA076292).

