## Supplementary Methods_Tables_Figures for "Ploidy shapes gemcitabine response through altered potency and delayed cell death"

### 2 Supplementary Information

#### 2.1 In-vitro Delay-aware cytostasis and cytotoxicity model for gemcitabine

##### 2.1.1 Intracellular dFdCTP signal surface

The default in vitro gemcitabine live/dead model is driven directly by a ploidy-specific intracellular dFdCTP signal surface rather than by parent extracellular gemcitabine pharmacokinetics. For ploidy $p \in \{2N, 4N\}$, administered gemcitabine dose $d$ (in $\mu$M), and time $t$ (in days), the PK-derived intracellular signal is denoted

$$C^{\mathrm{PK}}_{\mathrm{dFdCTP},p}(t, d).$$

This signal surface is built from the `dFdCTP (ng/mL)` PKPD measurements after conversion to $\mu$M-equivalent units and subtraction of the baseline ($t = 0$) signal for each calibrated PK sheet. Each calibrated dose retained its own measured time course. Terminal behavior was represented by a fitted exponential tail when the post-peak decay phase was identifiable; otherwise, a predefined fallback half-life of 1 day was used for that dFdCTP profile. Within the calibrated dose range, the surface interpolates dFdCTP values across dose in log-dose space. Below the minimum calibrated PK dose, the temporal profile at the minimum calibrated dose is preserved and scaled proportionally with dose. At zero administered dose, the intracellular dFdCTP signal is defined to be zero. Above the maximum calibrated PK dose, the default implementation uses explicit linear extrapolation in log-dose space based on the calibrated dose profiles evaluated at time $t$.

##### 2.1.2 Effective dose-response correction

For the default fits reported here, we used the beta–Hill baseline/confluence-death model. The effective intracellular dFdCTP signal combines a ploidy-specific power-law dose correction with a shared normalized Hill dose gate,

$$\widetilde{C}_p(t, d) = C^{\mathrm{PK}}_{\mathrm{dFdCTP},p}(t, d) \left(\frac{d}{d_{\mathrm{ref},p}}\right)^{\beta_p - 1} G(d; EC_{50}, h, d_{\mathrm{ref},p}),$$

where $d_{\mathrm{ref},p}$ is the minimum calibrated PK reference dose for ploidy $p$ and $\beta_p$ is a ploidy-specific dose-scaling exponent. This choice preserves the PK-derived signal when $\beta_p = 1$ and allows sublinear or supralinear effective dose scaling when $\beta_p \neq 1$. The shared Hill gate is

$$G(d; EC_{50}, h, d_{\mathrm{ref},p}) = \frac{H(d; EC_{50}, h)}{H(d_{\mathrm{ref},p}; EC_{50}, h)}, \qquad H(d; EC_{50}, h) = \frac{d^h}{EC_{50}^h + d^h}, \tag{2}$$

where $EC_{50}$ and $h$ are shared fit parameters. At $d = 0$, the effective signal is set to $\widetilde{C}_p(t, 0) = 0$.

##### 2.1.3 Immediate cytostasis and distributed delay to death

Drug-induced proliferation suppression is driven by the *immediate* effective dFdCTP signal. The cytostatic multiplier is

$$m_{\mathrm{cyto},p}(t, d) = \frac{1}{1 + k_{\mathrm{cyto},p}\widetilde{C}_p(t, d)},$$

where $k_{\mathrm{cyto},p}$ is a ploidy-specific cytostatic potency parameter and $m_{\mathrm{cyto},p} \in (0, 1]$.

Drug-induced death is delayed through a linear transit chain of length $n$:

$$\frac{dZ_{1,p}}{dt} = k_{\mathrm{tr},p}\left(\widetilde{C}_p(t, d) - Z_{1,p}\right),$$

$$\frac{dZ_{i,p}}{dt} = k_{\mathrm{tr},p}\left(Z_{i-1,p} - Z_{i,p}\right), \qquad i = 2, \ldots, n.$$

Here $k_{\mathrm{tr},p}$ is a ploidy-specific transit rate and $Z_{n,p}(t)$ is the delayed effective signal used for death. This linear chain provides a gamma-like distributed delay from intracellular dFdCTP exposure to death commitment. The number of transit compartments

$n$ is treated as a structural choice and selected by comparing fits over a small grid of integer candidate values, rather than estimated as a continuous parameter. For each replicate trajectory, the transit-chain states are initialized at zero, $Z_{i,p,r,d}(0) = 0$ for $i = 1, \dots, n$. In the default analysis, the candidate grid was $n \in \{2, 3, 4, 5, 6, 7\}$.

###### 2.1.4 Alive/dead cell-count dynamics

Let $A_p(t)$ denote the live-cell compartment and $D_p(t)$ the observed dead-cell compartment for ploidy $p$. The default model includes a ploidy-specific baseline death hazard $\mu_{\text{base},p}$ and a ploidy-specific drug-induced death term

$$\kappa_{\text{Gem},p}(t, d) = k_{\text{kill},p} Z_{n,p}(t),$$

where $k_{\text{kill},p}$ is the cytotoxic potency per unit delayed effective signal. Following the manuscript convention,

$$\Phi_{\text{Gem},p}(t, d) = \kappa_{\text{Gem},p}(t, d).$$

The default model also includes confluence-associated death through

$$\mu_{\text{ctrl},p}(A_p) = \mu_{\text{base},p} + \mu_{\text{conf},p} \left( \frac{A_p}{K_p} \right)^q,$$

with fixed exponent $q = 4$. The total death hazard is therefore

$$\lambda_p(t, d) = \mu_{\text{ctrl},p}(A_p(t)) + \kappa_{\text{Gem},p}(t, d).$$

The corresponding live/dead dynamics are

$$\frac{dA_p}{dt} = r_p \, m_{\text{cyto},p}(t, d) \, A_p \left( 1 - \frac{A_p}{K_p} \right) - \lambda_p(t, d) A_p,$$

$$\frac{dD_p}{dt} = \lambda_p(t, d) A_p - k_{\text{clear},p} D_p.$$

Here $r_p$ is the ploidy-specific maximum proliferation rate, $K_p$ is the ploidy-specific carrying capacity, and $k_{\text{clear},p}$ is the ploidy-specific disappearance or clearance rate of observed dead objects.

###### 2.1.5 Replicate-specific initial conditions

Each well/replicate trajectory was initialized from its own first paired alive/dead observation. Specifically, for replicate $r$, ploidy $p$, and dose $d$, the model initial conditions were set to

$$A_{p,r,d}(0) = A_{\text{obs},p,r,d}(t_0), \qquad D_{p,r,d}(0) = D_{\text{obs},p,r,d}(t_0),$$

where $t_0$ denotes the first retained imaging time point. Thus, replicate-to-replicate differences in seeding density and initial dead-object count were carried forward directly through the trajectory-specific initial conditions. Only time points with paired finite alive and dead observations were retained for likelihood evaluation.

###### 2.1.6 Ploidy-specific partial pooling and calibration.

The primary fit uses penalized maximum a posteriori partial pooling across ploidies in log-parameter space. For each positive ploidy-specific parameter $\alpha_p$,

$$\log \alpha_p = \mu_\alpha + \delta_{\alpha,p},$$

where $\mu_\alpha$ is a shared population-level log parameter and $\delta_{\alpha,p}$ is a ploidy-specific deviation penalized by a zero-centered Gaussian prior. In the default model, the ploidy-specific fitted parameters are

$$r_p, \; K_p, \; k_{\text{tr},p}, \; k_{\text{kill},p}, \; k_{\text{clear},p}, \; k_{\text{cyto},p}, \; \beta_p, \; \mu_{\text{base},p}, \; \mu_{\text{conf},p}.$$

The default model also fits the shared Hill-gate parameters $(EC_{50}, h)$. Simpler nested configurations, obtained by fixing $\beta_p$, disabling the Hill gate, or disabling confluence-associated death, were retained for sensitivity analyses. The corresponding prior penalty added to the objective is

$$\mathcal{P}_{\text{prior}} = \frac{1}{2}\sum_{\alpha}\sum_{p}\left(\frac{\delta_{\alpha,p}}{\sigma_\alpha}\right)^2,$$

with prior standard deviations $\sigma_r = \sigma_K = \sigma_{EC_{50}} = \sigma_h = 0.75$, $\sigma_{k_{\text{tr}}} = \sigma_{k_{\text{kill}}} = \sigma_{k_{\text{clear}}} = \sigma_{k_{\text{cyto}}} = \sigma_{\mu_{\text{base}}} = \sigma_{\mu_{\text{conf}}} = 1.00$, and $\sigma_\beta = 0.50$.

##### 2.1.7 Optimization routine

For model calibration, the alive-cell channel included only objects classified as Alive, whereas the dead-cell channel included objects classified as either Dead or Transitional. The calibration data consist of time-resolved live and dead object counts from IncuCyte assays across gemcitabine doses, replicates, and ploidies. Time is measured in days and administered gemcitabine dose in $\mu$M. The primary objective is a negative-binomial count likelihood evaluated on both the alive and dead channels. For an observed count $y$ with model mean $\mu$ and dispersion $\theta$,

$$Y \sim \text{NB}(\mu, \theta), \qquad \text{Var}(Y) = \mu + \frac{\mu^2}{\theta}.$$

Let $\theta_A$ and $\theta_D$ denote the dispersion parameters for alive and dead observations, respectively. The total objective minimized in the primary fit is

$$\mathcal{L} = -\sum_{\text{obs}} \log p_{\text{NB}}(A_{\text{obs}} \mid A_{\text{model}}, \theta_A) - \sum_{\text{obs}} \log p_{\text{NB}}(D_{\text{obs}} \mid D_{\text{model}}, \theta_D) + \mathcal{P}_{\text{prior}},$$

where $\mathcal{P}_{\text{prior}}$ is the Gaussian penalty associated with the ploidy-deviation terms in log space. Positive parameters are optimized on the log scale. Candidate transit-chain lengths $n$ are compared over a fixed integer grid, and the selected model is the one with the lowest raw posterior objective (data negative log-likelihood plus prior penalty). Optimization is performed with L-BFGS-B on an internally normalized objective for numerical conditioning, while raw negative log-likelihood and posterior objective values are retained for reporting and model comparison. ODE trajectories are solved with LSODA through `solve_ivp`. In the default configuration, the fitted positive parameters therefore comprise the ploidy-specific growth, delay, kill, clearance, cytostasis, dose-scaling, baseline-death, and confluence-death parameters together with the shared Hill-gate parameters $EC_{50}$ and $h$.

For calibration, live and dead counts were assembled into replicate-specific trajectories by matching observations across time, ploidy, gemcitabine dose, and well identity. The default analysis used the first 5 days of imaging data. Each trajectory was initialized from its own first paired alive/dead observation, and non-finite or unpaired alive/dead time points were excluded before likelihood evaluation. In the default phenotype aggregation, Alive objects defined the live channel, whereas Dead and Transitional objects together defined the dead channel.

**Supplementary Table 1.** Best-fit parameters for the in-vitro gemcitabine live/dead model.

| Parameter | Units | Brief definition | 2N | 4N |
|---|---|---|---|---|
| $r$ | day$^{-1}$ | Max. proliferation rate of live cells | 1.25 | 1.78 |
| $K$ | cells (or live objects) | Carrying capacity | 25,358 | 20,512 |
| $k_{\text{tr}}$ | day$^{-1}$ | Transit rate governing delay from effective dFdCTP exposure to death signal | 6.84 | 2.82 |
| $n_{\text{tr}}$ | dimensionless | Number of transit compartments in the linear-chain delay model | 5 | 5 |
| $k_{\text{kill}}$ | day$^{-1}(\mu\text{M dFdCTP})^{-1}$ | Cytotoxic potency per unit delayed intracellular drug signal | 217.3 | 129.9 |
| $k_{\text{clear}}$ | day$^{-1}$ | Clearance/disappearance rate of observed dead cells | 0.152 | 0.062 |
| $k_{\text{cyto}}$ | $(\mu\text{M dFdCTP})^{-1}$ | Cytostatic potency controlling suppression of proliferation by intracellular drug | 8041 | 1163 |
| $\beta$ | dimensionless | Dose-scaling exponent: administered gemcitabine –> intracellular drug signal | 0.253 | 0.488 |
| $\mu_{\text{base}}$ | day$^{-1}$ | Baseline drug-independent death hazard | 0.117 | 0.017 |
| $\mu_{\text{conf}}$ | day$^{-1}$ | Confluence-associated death hazard scale | 0.051 | 0.054 |

**Supplementary Table 2.** Symbol table.

| LaTeX symbol | Implementation name | Unit | Brief definition | Best-fit/default value |
|---|---|---|---|---|
| $A_p(t)$ | alive state | cell/object count | Live-cell count state for ploidy $p$. | Replicate-specific $A_{0,\text{rep}}$; dynamic prediction. |
| $D_p(t)$ | dead state | cell/object count | Observed dead-cell count state for ploidy $p$. | Replicate-specific $D_{0,\text{rep}}$; dynamic prediction. |
| $C^{\text{PK}}_{\text{dFdCTP},p}(t,d)$ | `DfdctpSignalSurface` | $\mu$M-equivalent dFdCTP | PK-derived intracellular dFdCTP signal before beta/Hill correction. | Derived from PKPD workbook; not a fitted scalar. |
| $\widetilde{C}_p(t,d)$ | effective corrected dFdCTP signal | $\mu$M-equivalent effective dFdCTP | Effective dFdCTP signal after beta correction and Hill gate. | Computed from $C^{\text{PK}}_{\text{dFdCTP},p}$, $\beta_p$, $EC_{50}$, and $h$. |
| $d_{\text{ref},p}$ | `min_calibration_dose_uM` | $\mu$M | Minimum calibrated PK reference dose for normalization. | 2N: 0.1; 4N: 0.1. |
| $\beta_p$ | `beta_dose` | dimensionless | Ploidy-specific power-law dose-scaling exponent. | 2N: 0.2533; 4N: 0.4877. |
| $EC_{50}$ | `dose_gate_ec50_uM` | $\mu$M | Shared Hill-gate half-activation dose. | 0.018529 $\mu$M (18.529 nM). |
| $h$ | `dose_gate_hill` | dimensionless | Shared Hill coefficient for dose gate. | 2.6928. |
| $Z_{i,p}$ | transit compartments | $\mu$M-equivalent effective dFdCTP | Transit-chain state $i$ for delayed drug-death signal. | Dynamic state; not separately fitted. |
| $n$ | `n_tr` | compartment count | Number of transit compartments selected by model comparison. | 5. |
| $k_{\text{tr},p}$ | `k_tr` | day$^{-1}$ | Ploidy-specific transit-chain rate. | 2N: 6.8427; 4N: 2.8202. |
| $k_{\text{cyto},p}$ | `k_cyto` | ($\mu$M-equivalent dFdCTP)$^{-1}$ | Ploidy-specific cytostatic potency. | 2N: 8040.9143; 4N: 1163.1135. |
| $k_{\text{kill},p}$ | `k_kill` | day$^{-1}$($\mu$M-equivalent dFdCTP)$^{-1}$ | Ploidy-specific cytotoxic potency per delayed signal. | 2N: 217.2640; 4N: 129.8923. |
| $r_p$ | `r` | day$^{-1}$ | Ploidy-specific maximum proliferation rate. | 2N: 1.2464; 4N: 1.7809. |
| $K_p$ | `K` | cell/object count | Ploidy-specific carrying capacity. | 2N: 25,358.2284; 4N: 20,512.0674. |
| $k_{\text{clear},p}$ | `k_clear` | day$^{-1}$ | Ploidy-specific disappearance rate for observed dead objects. | 2N: 0.1518; 4N: 0.0616. |
| $\mu_{\text{base},p}$ | `mu_base_death` | day$^{-1}$ | Ploidy-specific drug-independent baseline death hazard. | 2N: 0.1168; 4N: 0.0174. |
| $\mu_{\text{conf},p}$ | `mu_confluence_death` | day$^{-1}$ | Confluence-associated death scale at $A_p/K_p = 1$. | 2N: 0.0514; 4N: 0.0541. |
| $q$ | `confluence_death_exponent` | dimensionless | Fixed exponent for confluence-associated death. | 4.0. |
| $\theta_A$ | `theta_alive` | count-dispersion parameter | Negative-binomial overdispersion for alive counts. | 6.9519. |
| $\theta_D$ | `theta_dead` | count-dispersion parameter | Negative-binomial overdispersion for dead counts. | 7.7583. |

**Supplementary Table 3.** Combined effective dose-correction factors in the best-fit model. Values are normalized to the 100 nM reference dose.

| Gemcitabine dose | 2N correction | 4N correction |
|---|---|---|
| 3.125 nM | 0.111 | 0.049 |
| 6.25 nM | 0.408 | 0.213 |
| 12.5 nM | 1.229 | 0.755 |
| 25 nM | 1.967 | 1.421 |
| 50 nM | 1.586 | 1.348 |
| 100 nM | 1.000 | 1.000 |
| 200 nM | 0.601 | 0.707 |
| 400 nM | 0.359 | 0.497 |
| 800 nM | 0.214 | 0.348 |

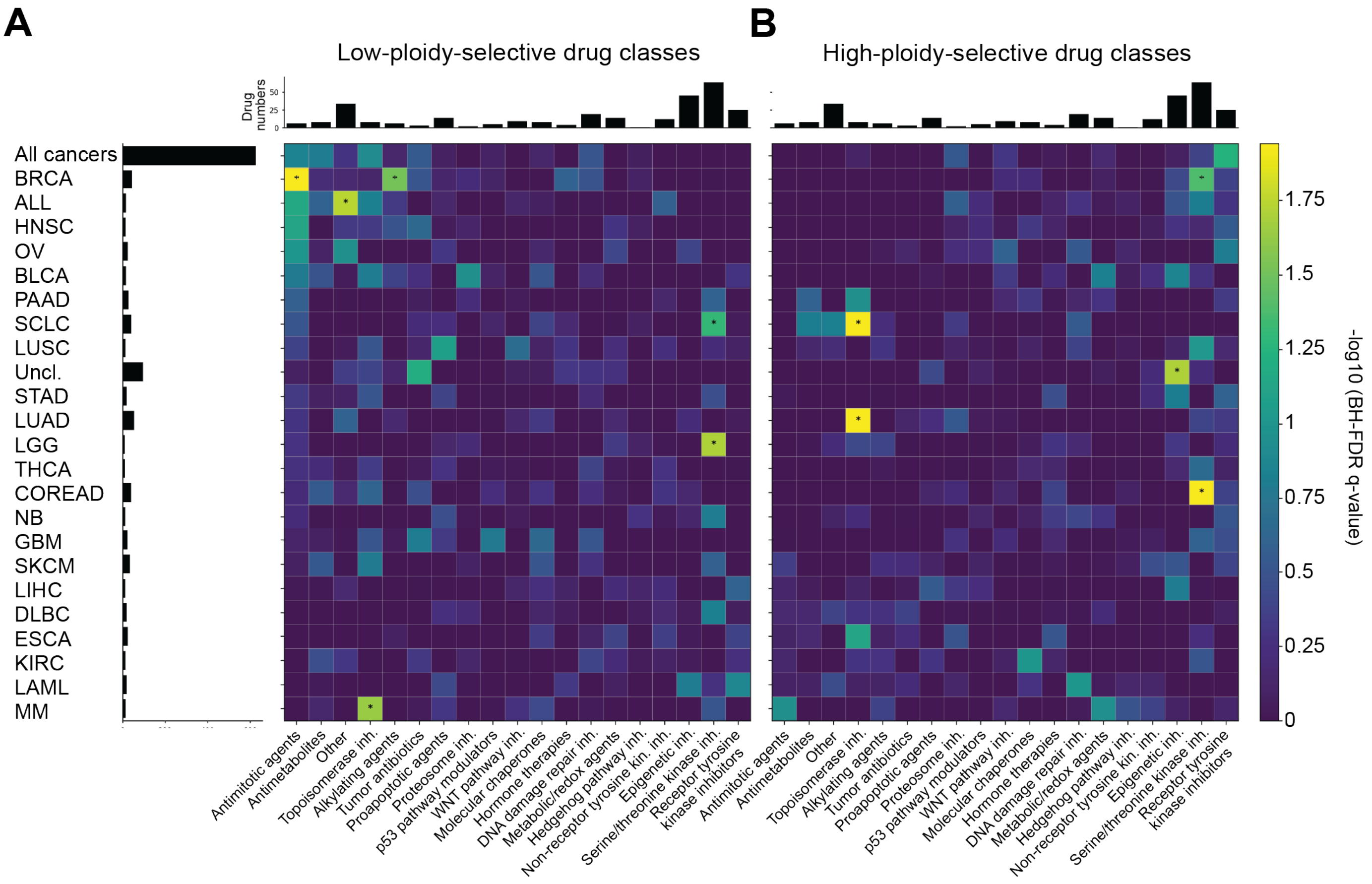


**Supplementary Figure 1. Expanded landscape of ploidy-associated drug-class enrichment across cancer cell lines.** (**A**) Heatmap showing drug classes enriched among compounds with response patterns consistent with preferential low-ploidy sensitivity. Rows indicate cancer types and columns indicate curated drug classes. Color represents enrichment significance as $-\log_{10}(p)$, with larger values indicating stronger enrichment. (**B**) Heatmap showing drug classes enriched among compounds with response patterns consistent with preferential high-ploidy sensitivity, using the same row order, column order, and color scale as in panel A. Stars indicate nominal enrichment significance at $p \leq 0.05$. Exact zero values from permutation testing were plotted at the empirical permutation floor to avoid infinite $-\log_{10}(p)$ values. (GDSC: Genomics of Drug Sensitivity in Cancer, AUC: Area Under Curve)

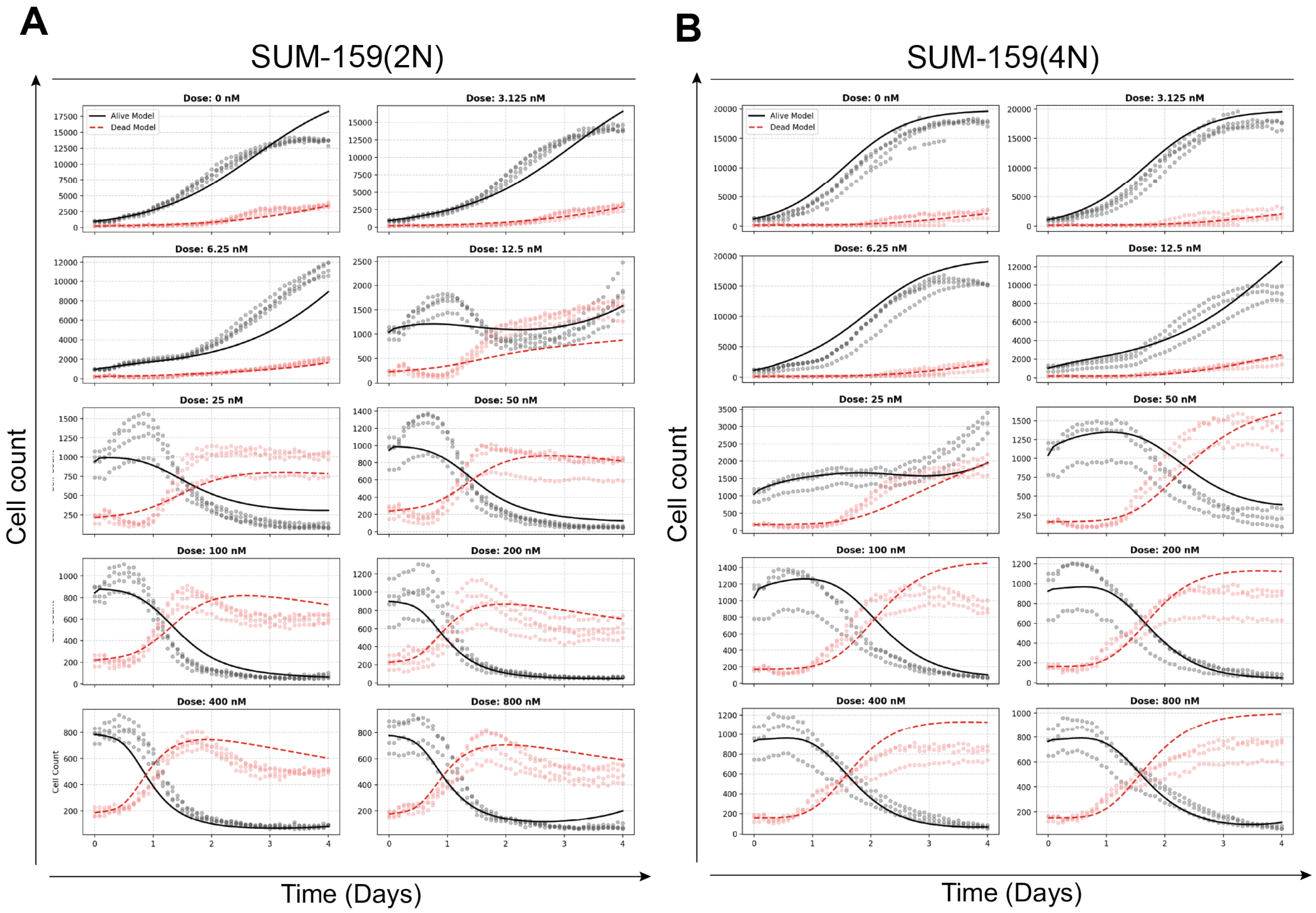


**Supplementary Figure 2. The delay-aware model recapitulates dose-dependent live and dead cell dynamics in both ploidy states.** (**A**) Joint cohort-level fits for near-diploid (2N) and (**B**) near-tetraploid (4N) SUM-159 cells. In each dose panel, points show observed alive and dead cells counts as objects over time across replicate wells. Solid and dashed curves show the corresponding model-predicted alive and dead trajectories from the best cohort-level fit. The fitted model was the intracellular dFdCTP-driven model, with immediate cytostasis, delayed drug-induced death, ploidy-specific $\beta$ dose correction, a fitted shared Hill dose gate, baseline death, and fitted confluence-associated death. Parameters were inferred jointly across doses within each ploidy using the alive/dead negative-binomial objective. Control (0 nM) trajectories are included to show baseline growth and death behavior in the absence of treatment.

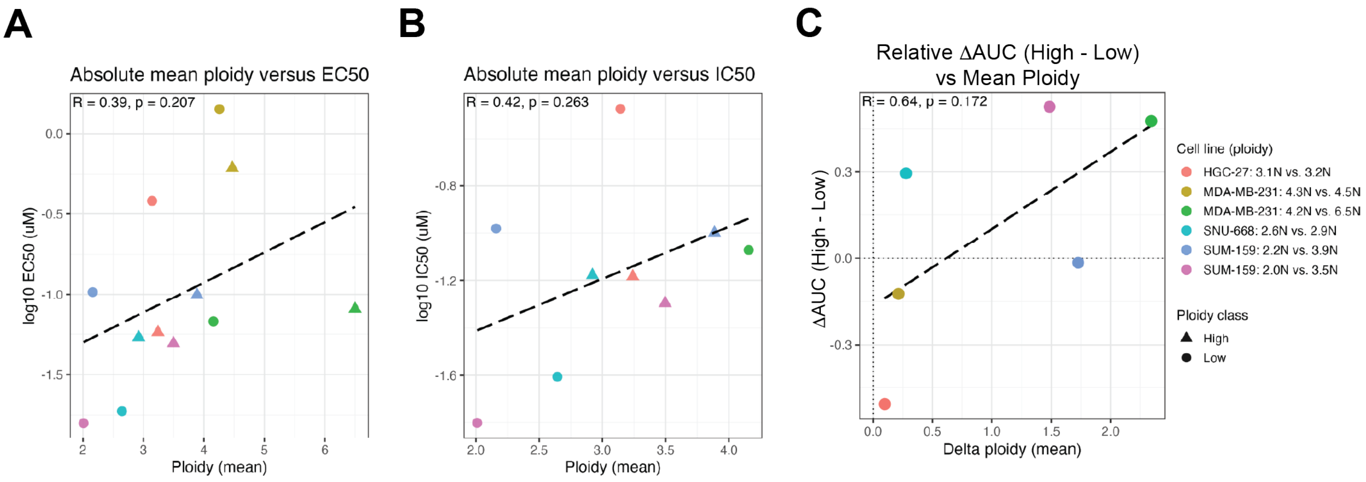


**Supplementary Figure 3. Complementary gemcitabine dose-response metrics support a directional association between increasing ploidy and gemcitabine resistance.** (**A**) Relationship between mean cell ploidy and fitted gemcitabine EC50 across individual samples. (**B**) Relationship between mean cell ploidy and fitted gemcitabine IC50 across samples for which IC50 could be estimated. (**C**) Paired comparison of the change in normalized gemcitabine dose-response AUC versus the change in mean ploidy between low- and high-ploidy isogenic lineage- pairs. Across these complementary metrics, higher ploidy showed directionally positive associations with reduced gemcitabine sensitivity, although the EC50, IC50, and paired delta-AUC associations did not reach statistical significance.

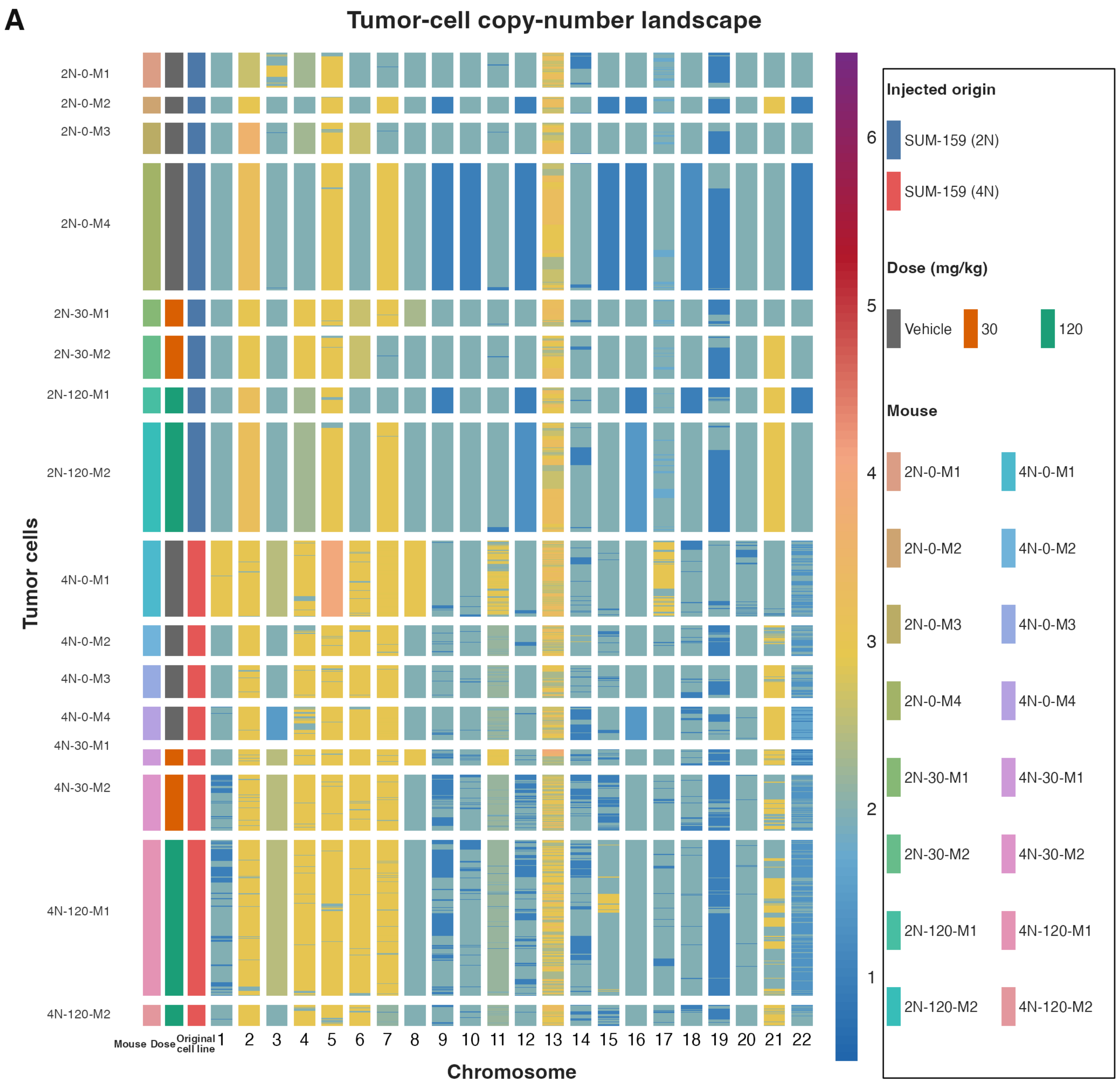


**Supplementary Figure 4. Tumor-cell chromosome copy number landscape.** **(A)** NUMBAT-derived chromosome-level copy-number profiles for the final 9,832 tumor cells selected from 14,125 cells with postprocessed NUMBAT estimates (4,880 from 2N-origin tumors and 4,952 from 4N-origin tumors; 5,335 gemcitabine-treated). Rows are cells, columns are chromosomes 1–22, and annotation bars indicate injected origin, treatment dose, and mouse.

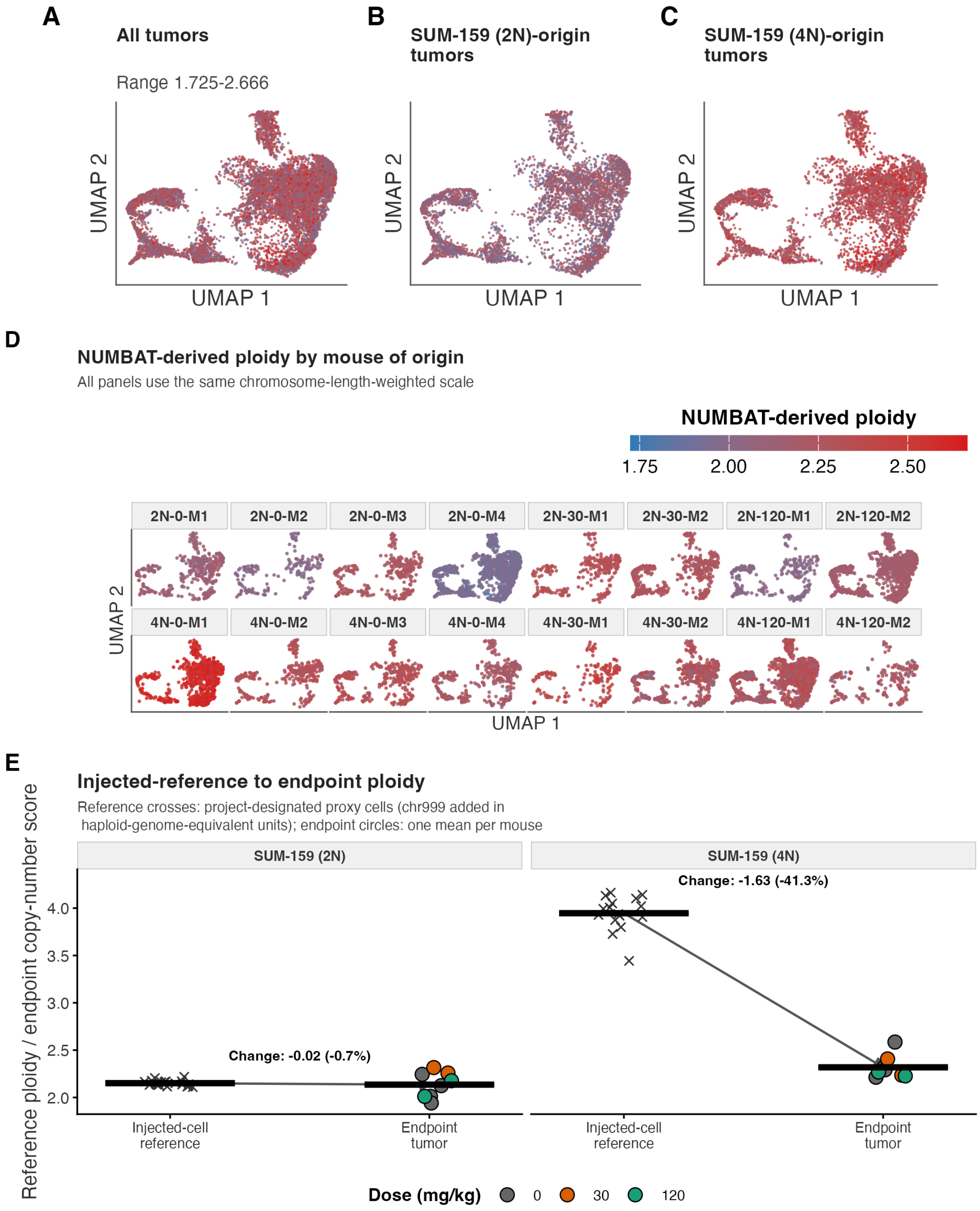


**Supplementary Figure 5. Endpoint tumor ploidy and NUMBAT-derived copy-number states across the xenograft-derived cellular landscape. (A–D)** NUMBAT-derived chromosome-length-weighted ploidy among the 9,832 final-QC tumor cells (4,880 from 2N-origin and 4,952 from 4N-origin tumors), displayed on the same 1.725–2.666 scale, with blue indicating lower and red higher ploidy. **(A)** All tumors combined. **(B,C)** Cells from 2N- and 4N-origin tumors, respectively. **(D)** The same cells faceted by mouse; facet headers identify mouse, injected origin, and dose. **(E)** Project-designated lineage-matched injected-state karyotype proxies (crosses; 20 2N-A7M and 16 4N-A5M metaphases) compared with one final-QC terminal mean per mouse (circles; eight mice per origin; color indicates dose). These proxy populations are not same-passage measurements of the A6M and A4M inocula. Reference ploidy is the sum of the autosomal chromosome-length-weighted estimate and the chromosome-999 unassigned-DNA value, in haploid-genome-equivalent units. Horizontal bars show the proxy-cell mean (2N, 2.151; 4N, 3.947) or the mean of the eight endpoint mouse means (2N-origin, 2.136; 4N-origin, 2.319), and arrows show the descriptive proxy-to-endpoint change. The 2N-origin mean changed by $-0.016$ ploidy units ($-0.7\%$), the 4N-origin mean was 41.3% below its reference proxy, and the 2N–4N mean separation contracted by 89.8%. No formal $P$ value is attached because each reference represents one culture-level biological unit, the endpoint tumors are independently replicated mice, and the two origin-specific NUMBAT outputs use different schemas and calibration.

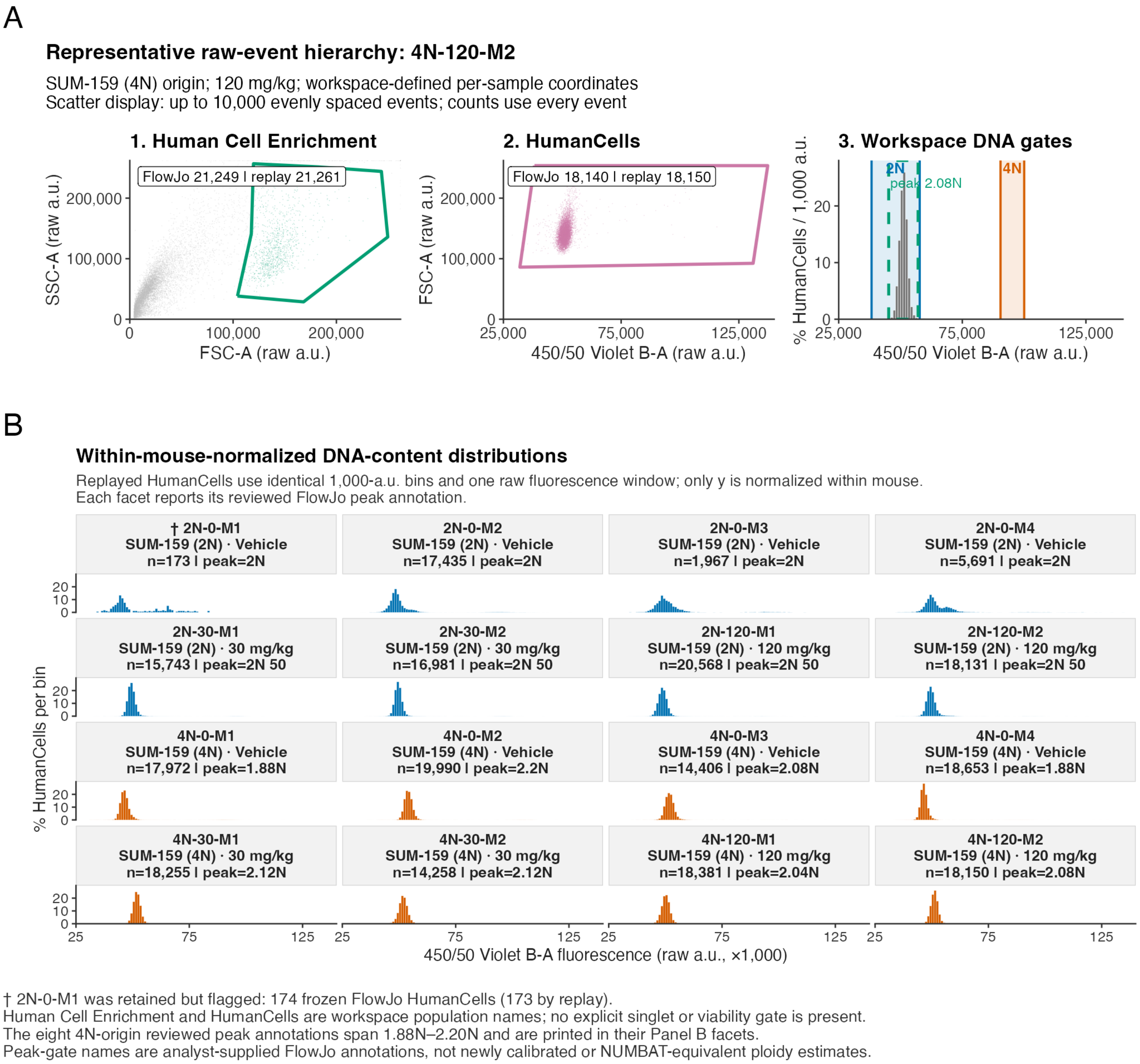


**Supplementary Figure 6. Endpoint tumor DNA-content flow cytometry.** **(A)** Representative raw-event gating hierarchy for 4N-A8-RR (120 mg/kg), selected as the QC-passing 4N-origin sample closest to the median frozen HumanCells count, with mouse ID used to break a tie. Human Cell Enrichment and HumanCells are workspace population names; the panel shows these sequential gates followed by the workspace DNA gates, and no explicit singlet or viability gate was present. Frozen FlowJo and full-precision raw-event replay counts are printed for the first two gates only. Scatter displays contain up to 10,000 evenly spaced events selected deterministically for legibility, whereas gate counts use all events. **(B)** Within-mouse-normalized DNA-content distributions for all 16 tumors (blue, 2N origin; orange, 4N origin), using identical 1,000-arbitrary-unit bins over the common raw 450/50 Violet B-A fluorescence window of 25,000–140,000 arbitrary units. Only the y-axis percentage is normalized within each HumanCells sample; fluorescence was not centered, peak-aligned, or rescaled. Each facet reports mouse, injected origin, dose, replayed HumanCells count, and the reviewed FlowJo peak annotation. The eight 4N-origin peak annotations span 1.88N–2.20N. The 2N-A1-0 sample was retained and marked despite its low HumanCells count (174 by frozen FlowJo; 173 by replay). Peak-gate names are analyst-supplied workspace annotations rather than newly calibrated or NUMBAT-equivalent ploidy estimates.

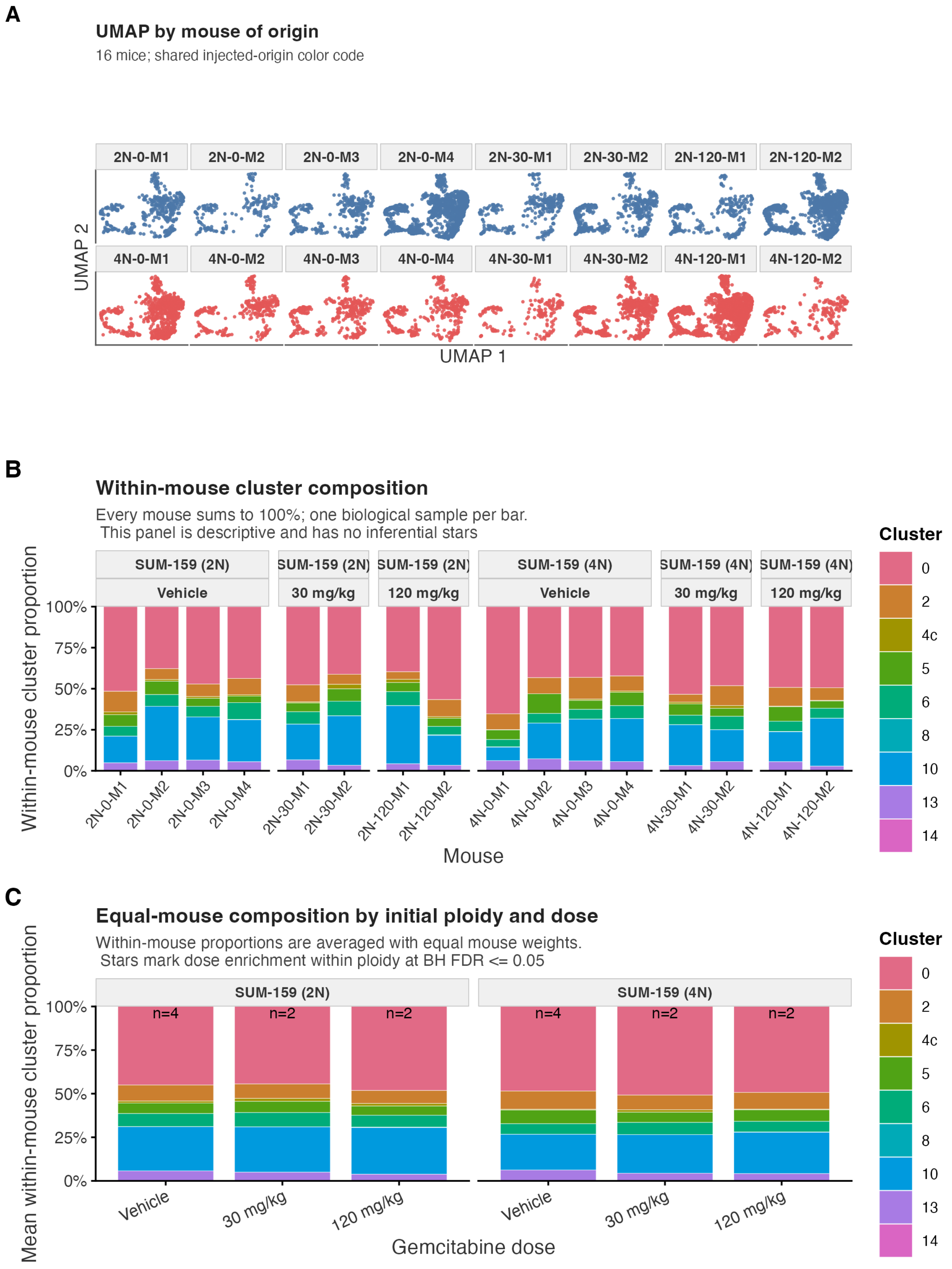


**Supplementary Figure 7. Mouse-resolved landscape and treatment-group composition of xenograft-derived tumor cells.** **(A)** UMAP of the 9,832 final-QC tumor cells from 16 mice, faceted by mouse; facet labels give injected origin and dose, and blue and vermilion identify 2N- and 4N-origin cells, respectively. **(B)** Mean within-mouse cluster proportions by injected origin and dose, with each mouse weighted equally; colors identify clusters and labels report the number of mice in each group. **(C)** For each cluster, positive enrichment of a displayed dose group relative to the remaining dose groups within the same injected-origin stratum was tested by one-sided exact permutation of dose labels on mouse-level proportions, with Benjamini–Hochberg correction across all dose-group-by-cluster contrasts. No contrast met $q \leq 0.05$ (minimum $q = 0.922$).

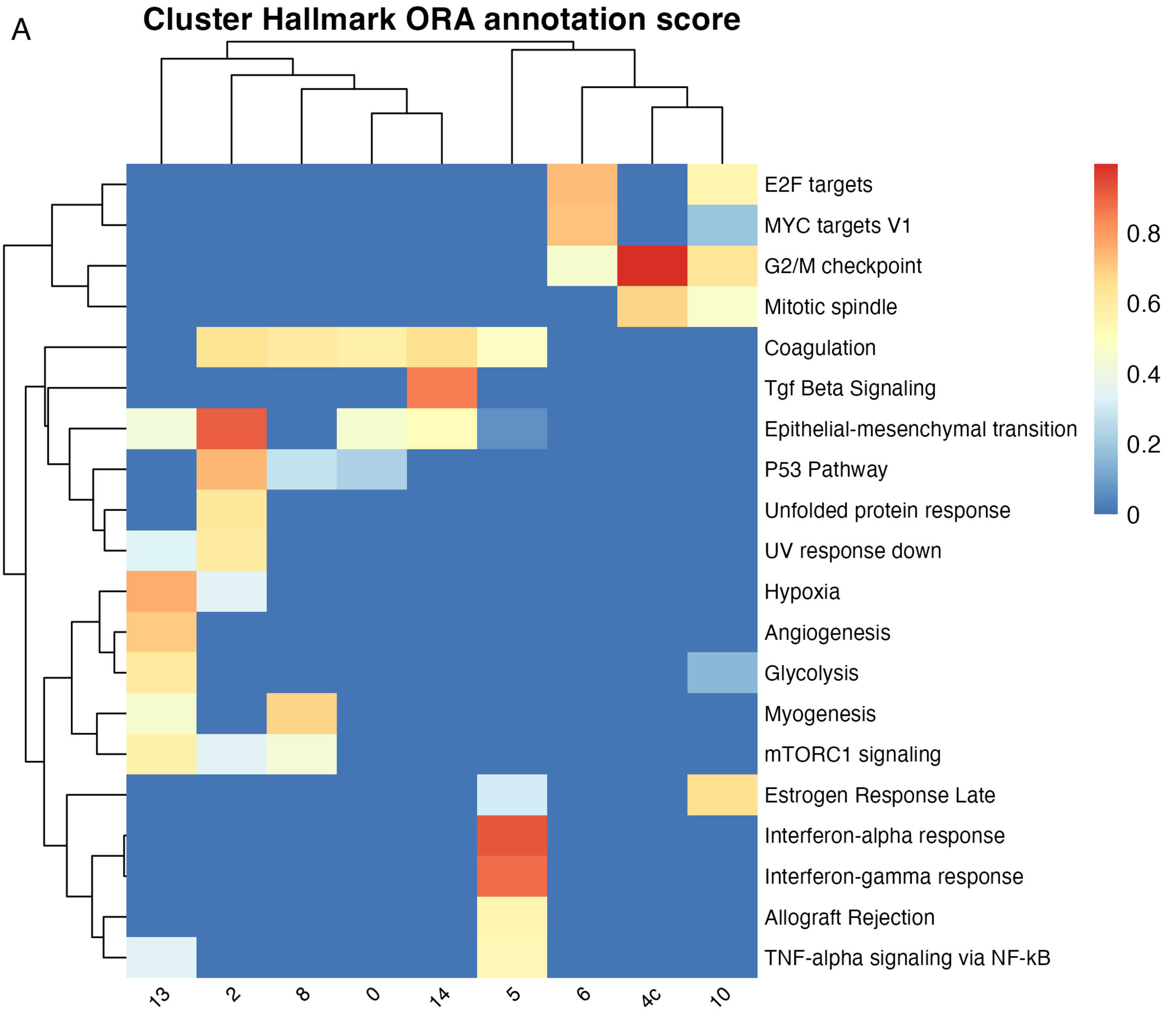


**Supplementary Figure 8. Cluster-level Hallmark over-representation analysis in the joint tumor–cell-line landscape. (A)** Cluster-versus-rest Hallmark over-representation analysis (ORA) across nine clusters in the joint 35,513-cell dataset (9,832 xenograft-derived tumor cells and 25,681 cultured SUM-159 reference cells). The analysis retained only GRCh38-assigned human features before normalization, differential-expression testing, and ORA; GRCm38-assigned mouse features were excluded. For each cluster, ORA used up to the top 100 significant, positively enriched markers (Benjamini–Hochberg-adjusted $P < 0.05$, absolute log fold-change $\geq 0.25$, and absolute detection-frequency difference $\geq 0.05$). One-sided hypergeometric ORA results were Benjamini–Hochberg corrected within cluster; retained terms had adjusted $P < 0.05$, at least three overlapping genes, and a Hallmark set size of 15–500 genes. Color shows a 0–1 annotation score, not a $P$ value: the mean of within-cluster min–max-scaled marker-effect, marker-detection, and overlap components. Rows are the 20 Hallmark sets with the largest maximum annotation score across clusters; cluster–pathway combinations without a positive retained score appear as 0. Hierarchical clustering changes row and column display order only.

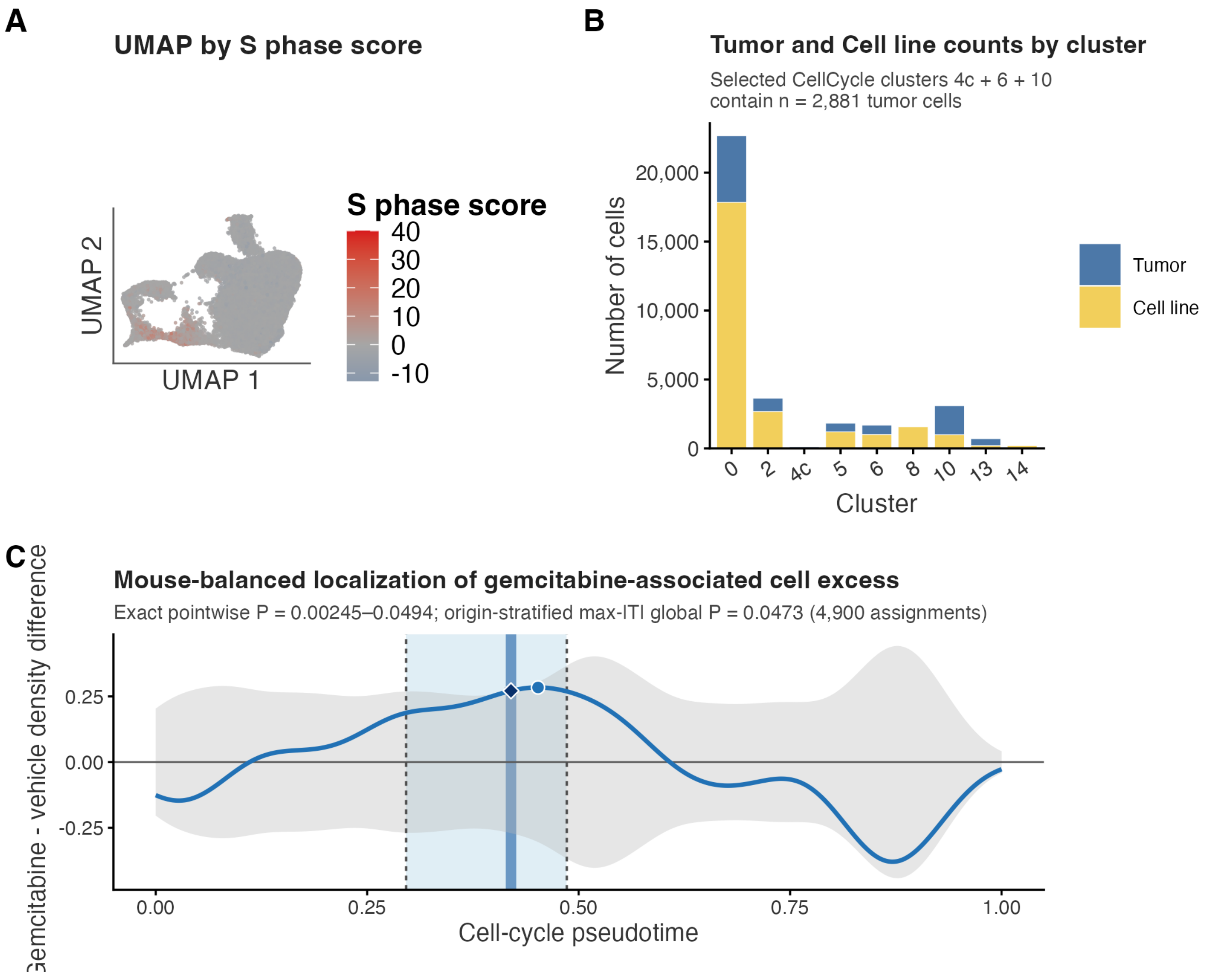


**Supplementary Figure 9. Cell-cycle state definition and gemcitabine-associated pseudotime localization.** **(A)** Joint UMAP of 35,513 cells (9,832 final-QC xenograft-derived tumor cells and 25,681 cultured SUM-159 reference cells), colored from blue through white to red with increasing Seurat S-phase score. **(B)** Raw cell counts by cluster and sample context (blue, tumor; yellow, cultured reference). Clusters 4c, 6, and 10 operationally define the cell-cycle-associated subset and contain 89, 686, and 2,106 tumor cells, respectively (2,881 total). **(C)** Mouse-balanced localization of the gemcitabine-associated cell excess along cell-cycle pseudotime among these 2,881 tumor cells from 16 mice (1,328 vehicle and 1,553 treated cells; eight mice per group; the treated group pools the 30- and 120-mg/kg doses). The curve is the equal-mouse treated-minus-vehicle Gaussian-kernel density contrast, and the gray band is its two-sided 95% simultaneous permutation null envelope. All 4,900 injected-origin-stratified assignments preserve four vehicle and four treated mice within each injected-origin stratum. The pale-blue band marks the maximal connected positive region with nominal two-sided exact pointwise permutation support (0.296–0.486; 96 grid points; pointwise $P = 0.00245$–0.0494); dashed lines mark these exact boundaries, which define the interval modeled in Fig. 7I. No single region-level $P$ value was calculated for this broad interval. The dark-blue band and curve segment mark the narrower 0.414–0.426 interval in which the observed contrast exceeded the studentized max-$|T|$ envelope; the corresponding global family-wise exact permutation test gave $P = 0.0473$. The circle marks the largest raw density difference at 0.452, and the diamond marks the strongest standardized evidence at 0.420.

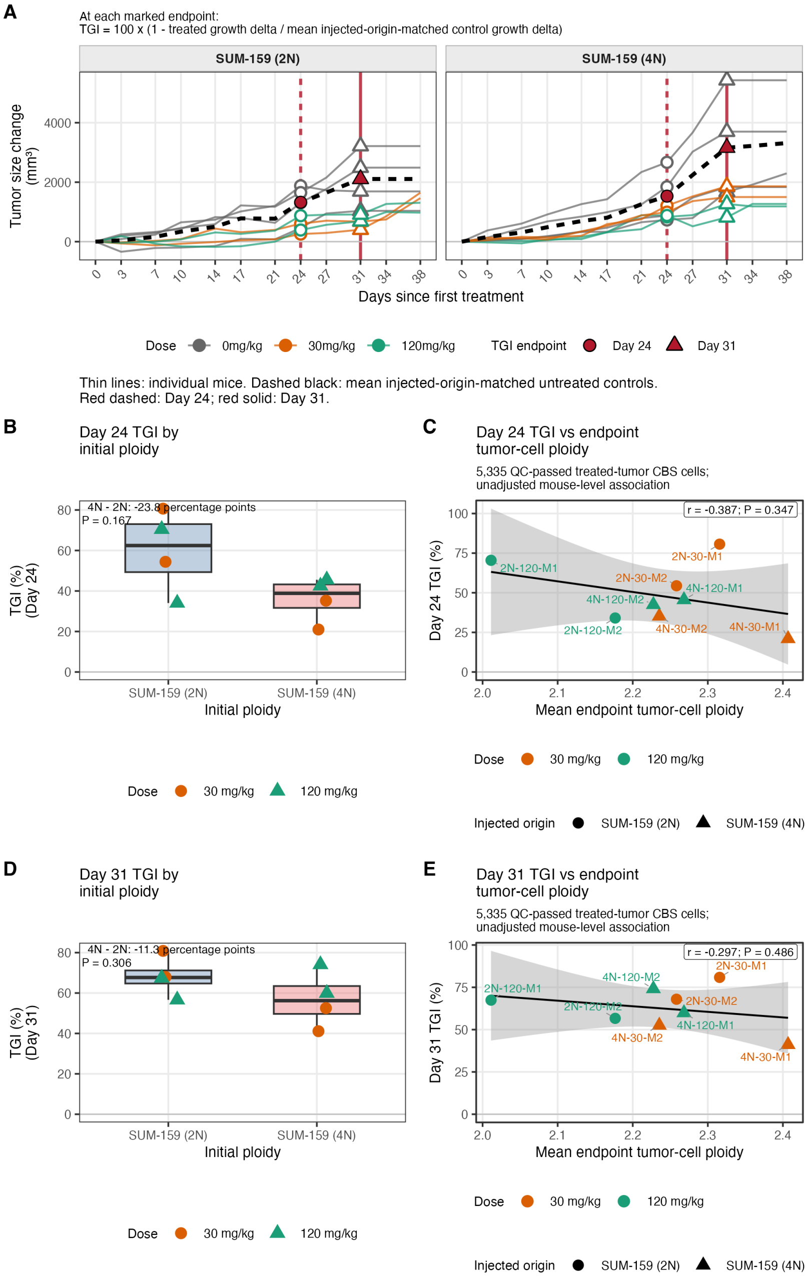


**Supplementary Figure 10. Later TGI endpoints preserve the direction but attenuate the magnitude of the injected-origin and endpoint-ploidy associations.** **(A)** Baseline-adjusted tumor-growth trajectories for tumors initiated with near-diploid (2N) or near-tetraploid (4N) SUM-159 cells, with Days 24 and 31 marked as alternative TGI endpoints. Thin lines show individual mice and dashed black lines show the mean untreated-control trajectory matched by injected origin. At each endpoint, TGI was calculated as $100 \times [1 - (\text{treated growth delta}/\text{mean matched-control growth delta})]$. **(B)** Day-24 TGI in treated tumors grouped by injected origin. The dose-adjusted 4N-minus-2N difference was -23.8 percentage points (exact dose-stratified permutation $P = 0.167$; $n = 4$ tumors per injected-origin group). **(C)** Unadjusted mouse-level association between Day-24 TGI and mean endpoint tumor-cell ploidy (Pearson $r = -0.387$, exact unrestricted permutation $P = 0.3469$; $n = 8$). **(D)** Day-31 TGI in treated tumors grouped by injected origin. The dose-adjusted 4N-minus-2N difference was -11.3 percentage points (exact dose-stratified permutation $P = 0.306$; $n = 4$ tumors per injected-origin group). **(E)** The corresponding unadjusted association at Day 31 (Pearson $r = -0.297$, exact unrestricted permutation $P = 0.4857$; $n = 8$). In B–E, only treated tumors are included; colors indicate dose, and shapes in C and E indicate injected origin. Lines and 95% confidence intervals in C and E show the raw mouse-level fits. Panels C and E use the same 5,335-cell QC-retained treated-tumor universe as Fig. 7L.
